# Intelligent Tool Orchestration for Rapid Mechanistic Model Prototyping: MCP Servers as AI-Biology Interfaces

**DOI:** 10.1101/2025.09.10.675105

**Authors:** Marco Ruscone, Miguel Vazquez, Alfonso Valencia

## Abstract

The construction of multicellular mechanistic models in systems biology typically requires months of literature research, programming expertise, and deep knowledge of specialized computational tools. Here we present intelligent tool orchestration through Model Context Protocol (MCP) servers that enable Large Language Models (LLMs) agents to function as AI laboratory assistants for rapid model prototyping. We demonstrate this approach by constructing a multiscale model of cancer cell fate in response to TNF, entirely through natural language interactions, using an AI agent connected to MCP servers that interface with three complementary modeling software: NeKo for constructing gene regulatory networks from prior-knowledge databases, MaBoSS for simulating and analyzing Boolean models, and PhysiCell for setting up multicellular agent-based models. Our architecture encompasses more than 60 specialized tools, covering the entire workflow from data collection to multicellular simulation setup. From this use case we derived three key design principles for biological AI-tool integration: first, optimal tool granularity is best defined at biological decision points, where domain knowledge guides modeling choices; second, comprehensive session management is essential for tracking complex workflows and ensuring reproducibility across long interactions; and third, effective orchestration must be established from the start to let LLMs combine tools flexibly while maintaining biological coherence. In applying this framework across three different LLMs and scenarios, we observed consistent end-to-end orchestration as well as model-dependent differences in network size, dynamical behaviors, and integration details. This variability highlights both the portability of our approach and the importance of careful reporting and validation when deploying different LLMs in scientific contexts. While the resulting models require refinement, this work establishes the foundation for AI-assisted rapid prototyping in systems biology, enabling researchers to explore computational hypotheses at unprecedented speed while maintaining biological fidelity.

## Introduction

Biological systems exhibit emergent behavior across multiple scales—from molecular interactions and gene regulatory networks to cellular dynamics and tissue-level phenomena. Understanding these multiscale processes is fundamental to biology, as cellular phenotypes emerge from complex interactions between genetic networks, signaling pathways, mechanical forces, and environmental cues(1–3). Multiscale mechanistic models have become essential tools for integrating this complexity, serving as “virtual laboratories” that enable researchers to test hypotheses, predict outcomes, and quickly explore untested scenarios.

Frameworks such as PhysiCell(4), PhysiBoSS(5), CompuCell3D(6), BioDynaMo(7), and Morpheus(8) have been developed to support such multiscale exploration, offering powerful virtual laboratories for biological hypothesis testing. The models simulated by the aforementioned softwares allow to study cancer cell differentiation, drug responses, and the effects of mechanical stimuli on cellular phenotypes(1, 9). However, the construction of such models presents significant barriers to entry for many researchers.

Traditional mechanistic modeling requires expertise across multiple domains: deep knowledge of biological pathways, proficiency in programming languages, familiarity with specialized software architectures, and understanding of complex mathematical frameworks(3). Researchers must navigate intricate workflows involving database queries, network construction algorithms, parameter estimation, and simulation software. This complexity has led to the development of automated pipelines (10) and graphical user interfaces (8, 11), yet these solutions still require substantial technical expertise and often lack the flexibility needed for novel research questions(12).

Most recently, human-interpretable grammar systems—as demonstrated in the paper by Johnson and colleagues(13)—have shown how natural language statements can be directly translated into mathematical models, representing a critical effort toward more accessible model construction.

The emergence of Large Language Models (LLMs) has opened new possibilities for scientific computing, offering natural language interfaces to complex computational tasks(14). However, integrating LLMs with specialized scientific software has remained challenging due to the need for structured, reliable connections between AI systems and domain-specific tools.

The key breakthrough in this integration challenge was tool calling (15, 16), which enables LLMs to invoke external capabilities in a structured way. Building on this, the Model Context Protocol (MCP) provides a standardized, REST-like framework for packaging tools and supporting secure, mostly remote use. MCP connects LLMs to external tools and data sources, enabling controlled, reliable interactions between AI assistants and scientific software. With MCP, LLMs can access vast knowledge bases, execute computational tasks, query databases, and generate results, while maintaining proper attribution and reproducibility.

For systems biology, this creates unprecedented opportunities for conversational modeling. Rather than requiring researchers to master complex software interfaces, MCP enables natural language interactions with sophisticated computational tools. The protocol allows LLMs to access vast biological knowledge while executing computational tasks, querying databases, and generating reproducible results. This represents a fundamental shift from code-centric to conversation-centric modeling, where AI assistants function as intelligent laboratory partners.

In this work, we present the first demonstration of intelligent tool orchestration for mechanistic modeling - an approach where AI assistants function as laboratory collaborators, rapidly prototyping complex computational models through natural language interactions. Following the digital twin approach, our objective is the full construction of a mechanistic multiscale model-from gene regulation to tissue dynamics-just by conversating with an LLM. To achieve this, we developed MCP servers for three established systems biology tools: NeKo, a python package for automatic gene regulatory network construction (17), MaBoSS, a C++ Boolean network simulator with a python interface (18), and PhysiCell (4), a multiscale agent-based modeling framework C++ based. Specifically, PhysiCell lends itself very well to this approach thanks to its interpretable human grammar feature (13), which uses a dictionary of signals (cellular stimuli) and behaviors (cellular phenotypes) to generate hypotheses using a natural language processing approach. Our ecosystem encompasses more than 60 specialized biological tools, supporting the complex tasks that span from data collection to model construction. These tools are organized according to novel architectural principles including optimal tool granularity, comprehensive session management, and intelligent workflow orchestration designed specifically for biological AI-tool integration. The seamless integration of these tools leads each scale feeding naturally into the next: molecular networks → gene regulation → cellular behavior → tissue dynamics → population emergent properties.

Through a case study of TNF response modeling, we demonstrate how tools-enabled AI assistants can exhibit sophisticated biological reasoning, automatically selecting appropriate molecular components, cell types, and parameters leveraging curated resources. The complete multiscale model—from molecular networks through Boolean dynamics to tissue simulation—was constructed in minutes rather than the months typically required for traditional model development, while maintaining biological plausibility and incorporating biologically-motivated parameter estimates where available in the literature.

Our approach represents a fundamental shift from code-centric to assistant-centric modeling, where AI systems serve as intelligent laboratory partners that can rapidly prototype models for hypothesis exploration (19). This work establish-esthe fundational architected of AI-assited scientific modeling and a first demonstration of the capacity of this technology to ensamble a working system that can be integrated in a larger framework for real work operations, including data query, autonomous model construction, validation, and optimization.

To assess portability and robustness of assistant-centric modeling, we explored how three differents LLMs performs model construction and analysis across three scenarios - TNF multiscale simulation construction, PhysiCell rule synthesis, and MaBoSS model refinement - with identical prompts and toolchains. We observed reliable orchestration across models with distinct trade-offs in mechanistic depth and outcome diversity, motivating design choices that accommodate LLM diversity and reinforcing the need for careful validation workflows.

## The Conversational Modeling Paradigm

The convergence of LLMs with domain-specific scientific tools represents a fundamental shift in how computational biology can be practiced. Rather than requiring researchers to master programming languages and complex software interfaces, conversational modeling enables direct dialogue between domain experts and computational tools through natural language interactions. It is therefore important to understand how this new technology works and how it can be exploited.

MCP servers act as a central hub that exposes different AI tools and data sources so they can work together in a secure and standardized way. Tools are the individual computational functions that the AI can invoke. Each tool represents a specific scientific operation with well-defined inputs, outputs, and biological meaning. Once the server is installed, the language model gains access to all the tools it contains. An ensemble of metadata embedded in the servers, helps the model understand how to use each tool and orchestrate them together. As shown in Fig 1, panel A, this architecture places the LLM agent at the top, mediating natural language requests and orchestrating workflows through the MCP layer, which in turn provides access to the specialized modeling servers.

**Fig. 1.**
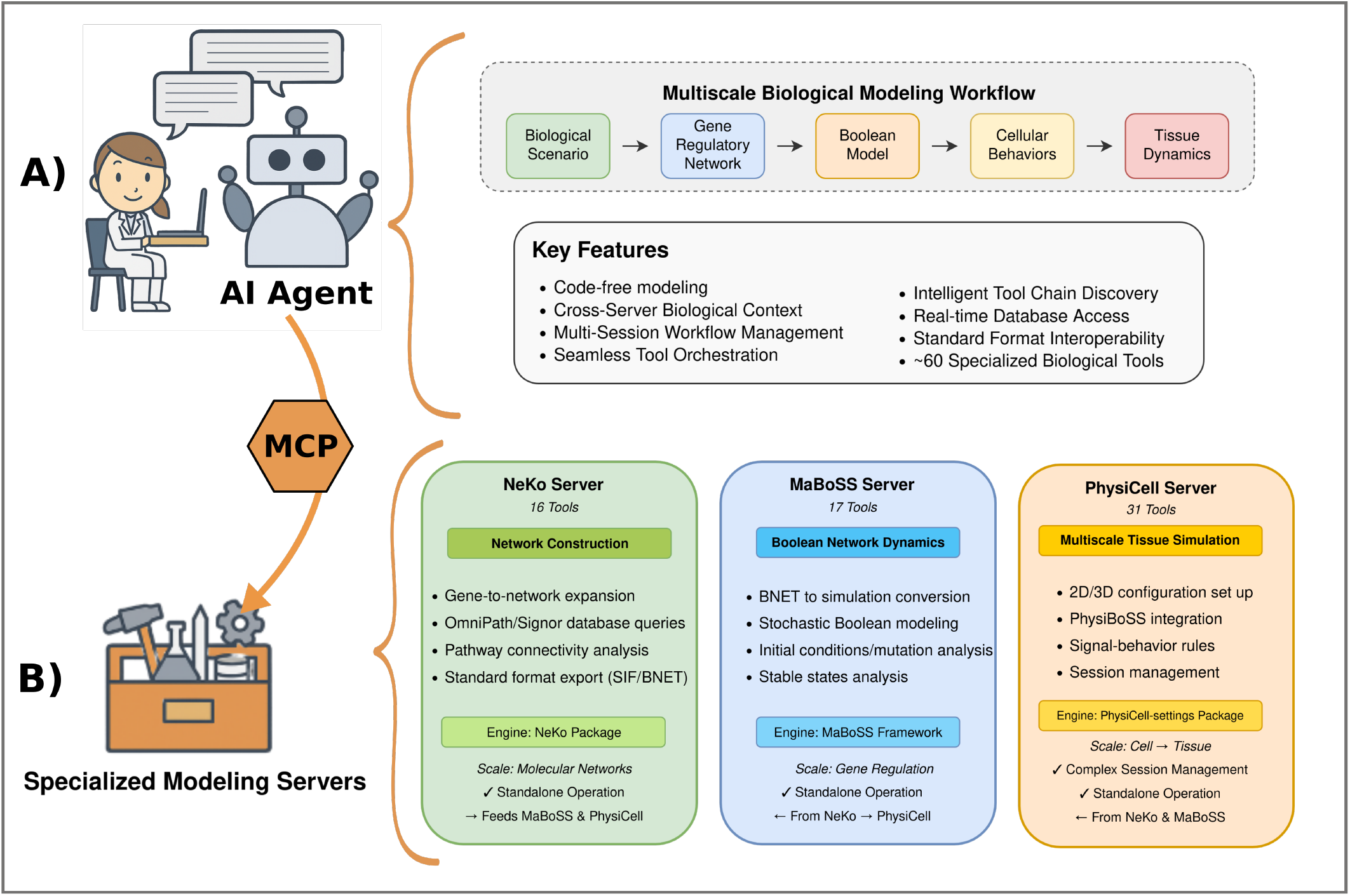
Conversational modeling ecosystem enabled by MCP servers. **A): Conceptual overview**. An LLM agent mediates natural language requests and orchestrates workflows through the Model Context Protocol (MCP), which provides standardized access to specialized modeling servers. This design allows domain experts to interact with computational tools via conversation rather than code. **B): Servers implementation**. Three MCP servers—NeKo, MaBoSS, and PhysiCell—collectively provide over 60 tools spanning molecular network construction, Boolean model dynamics, and multiscale tissue simulation. Together, they form a pipeline from molecular to tissue-level modeling, illustrating how conversational interfaces can unify the full mechanistic modeling workflow.

### A. From Code to Conversation

Traditional mechanistic modeling follows a code-centric workflow: researchers write scripts to query databases, construct networks, implement mathematical models, and analyze results. Each step requires specific programming knowledge and tool expertise. Conversational modeling inverts this relationship—researchers express their biological questions and modeling intentions in natural language, while AI assistants orchestrate the underlying computational tools.

This transformation affects the entire modeling ecosystem. Network construction shifts from manual database queries and scripting to guided conversations about pathway biology. Dynamic modeling transitions from writing mathematical equations to describing biological relationships and regulatory logic. Multiscale integration evolves from complex simulation programming to discussions about cellular behaviors and tissue-level phenomena.

### B. Architectural Principles

Our implementation demonstrates this paradigm through three complementary MCP servers (Fig 1, panel B) that collectively provide more than 60 specialized tools spanning the complete mechanistic modeling workflow. The architecture follows a layered design with natural language interface capabilities at the top, standardized tool interface protocols in the middle, and specialized computational biology servers at the foundation.

The **NeKo MCP server** (16 tools) exploits the NeKo python package to consruct molecular network from prior knowledge databases including OmniPath (20) and SIGNOR (21), providing gene-to-network expansion, pathway connectivity analysis, and standard format export capabilities (SIF and BNET). The **MaBoSS MCP server** (17 tools) is based on the MaBoSS framework, which simulates Boolean models by applying stochastic markoc processes on Boolean network. This server provides with tools to simulate analyze and simulate a Boolean model, changing the initial conditions, performing mutations and providing attractors (network stable states) analysis. The **PhysiCell MCP server** (28 tools) leverages the PhysiCell-config python package (https://github.com/marcorusc/PhysiCell_Settings) allowing to draft multiscale tissue simulation with 3D modeling capabilities. It includes MaBoSS integration (PhysiBoSS), drafting of cellular rules through Physicell’s human interpretable grammar and complex session management.

The key architectural insight is that effective conversational modeling requires more than simple API wrapping. The servers implement domain-specific language processing, cross-server biological context management, multi-session workflow orchestration, and biological validation guidance. Each server maintains standalone operation capability while supporting seamless data exchange through standardized formats (SIF, BNET), creating a multiscale pipeline from molecular networks through gene regulation to tissue dynamics.

### C. Novel Design Principles for Biological AI-Tool Integration

Designing MCP servers for biological modeling is more than wrapping APIs: tools that are too fine-grained can confuse the LLM (leading to wrong tools calls), while tools that are too coarse lock entire workflows into single calls. The challenge is to balance granularity and guidance so the model can reason about biology, chain tools effectively, and recover from errors without losing flexibility. Through extensive experimentation with different LLM models and architectural patterns, we designed key principles that distinguish effective biological tool orchestration from simple API wrapping:

#### Optimal Tool Granularity

As stated in different MCP developing guidelines from Anthropic https://modelcontextprotocol.io/quickstart/server or Visual Studio Code https://code.visualstudio.com/api/extension-guides/ai/mcp, we found that high-level granularity enables better modularity and AI reasoning. Each tool is extensively documented with comprehensive docstrings and parameter explanations. Tools that are too granular (micro-operations) prevent the AI from effective reasoning, while tools that are too coarse (complex workflows) limit flexibility. We achieved an optimal granularity by designing tools around workflow steps where biological knowledge is needed—for example, choosing the right parameters for constructing a network, deciding on relevant mutations to simulate, or defining a cell type and its cell cycle model. This allows the AI to reason about the biology rather than micromanaging technical operations.

#### Comprehensive Session Management

Complex biological workflows require state tracking across multiple tool calls. For the PhysiCell server, we implemented a session manager with unique identifiers that maintains workflow history and completion percentage tracking. This prevents incompatible XML structures and enables workflow resumption, as incomplete configurations cannot interface with standard PhysiCell inputs.

#### Intelligent Error Handling and Tool Chaining

Robust error messages with biological context trigger are injected in the LLM to enable tool chaining responses. Rather than generic API failures, errors include biological interpretation and suggest the LLM next steps to take, enabling the AI to adaptively modify workflows based on biological constraints.

#### Structured Output with Biological Context

Each tool provides clear documentation of operations performed and biological significance of results. This enables the AI to maintain biological reasoning chains across tool calls and make informed decisions about subsequent steps.

These principles were derived from iterative testing across different LLMs, ranging from lightweight models that required explicit guidance to more advanced models capable of autonomous biological reasoning. We found that maintaining optimal tool granularity at biological decision points, explicit session/state management, and structured, self-describing outputs are pivotal for robust orchestration across different LLMs. Together, these elements help to reduce prompt sensitivity, support tool chain recovery during the workflow orchestration, and can support reproducibility despite differences in the underlying language model.

## Proof of Concept: Demonstration of Work-flow Orchestration for Mechanistic Modeling

We have explored how three different LLMs (*GPT-5, GPT o4 mini, Sonnet-4*) adapt and respond to three different scenarios using identical prompts and the same MCP toolchain (NeKo, MaBoSS, PhysiCell). These scenarios can be addressed through two complementary interaction modes: a one-shot orchestration approach, where a single high-level prompt triggers the entire workflow autonomously (Fig. 2, Panel A), and an interactive conversation, where the user iteratively guides the LLM by selecting tools and parameters Fig. 2, Panel B). In this work, we focused on the autonomous orchestration mode, to assess the ability of each LLM to manage the workflow and autonomously call the appropriate tools, thus injecting a single prompt without further conversation. For each scenario we tested how the MCP servers operated both in combination and stand-alone mode, and then compared the outputs produced by the different LLMs: the multicellular mechanistic models generated in scenario 1 (Fig 3 Panel B), the sets of cell rules inferred in scenario 2 (Fig 3 Panel C), and the Boolean network attractors obtained in scenario 3 (Fig 3 Panel D). It is important to mention that at this stage our goal was not to assess the biological accuracy of the generated models, but rather to test whether the whole ecosystem works end-to-end. This is motivated by 2 choice: 1) none of the selected LLM (accessible through GitHub copilot), are specifically trained on biological information (which could in principle lead to more accurate models) and 2) our focus was on validating interoperability and tool orchestration rather than biological accuracy.

**Fig. 2.**
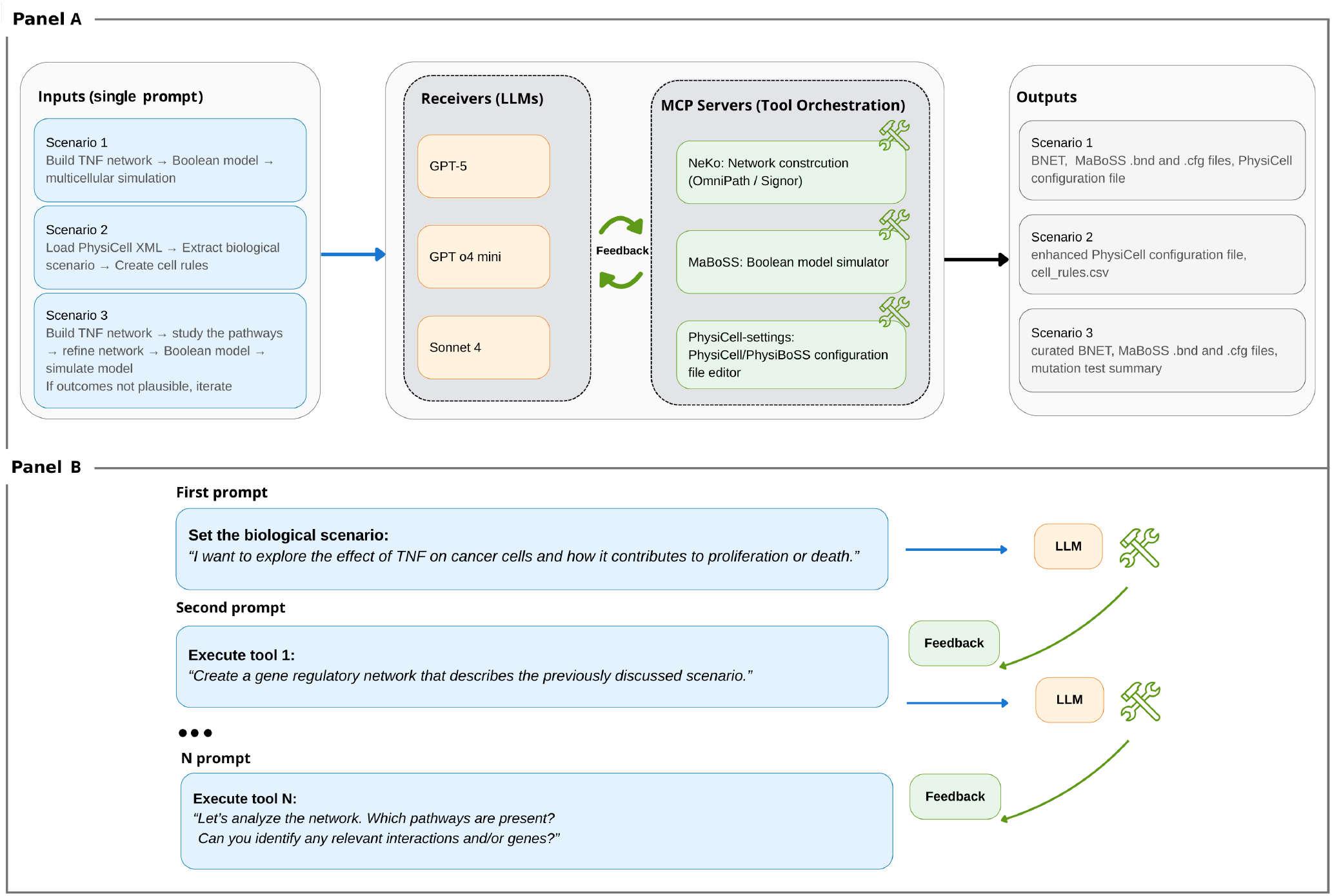
Approaches to conversational modeling with MCP servers. **Panel A: One-shot orchestration**. A single high-level prompt is provided to the LLM, which autonomously selects and sequences the appropriate tools. The model iterates internally with feedback from the MCP servers until a complete output is produced (e.g., a static network, a Boolean model, or a PhysiCell configuration files). **Panel B: Interactive conversation**. The workflow is guided step by step by the user, who provides successive prompts to decide which tools to execute and with which parameters. The LLM supports reasoning, tool invocation, and feedback handling, while the user retains fine-grained control of the modeling process.

**Fig. 3.**
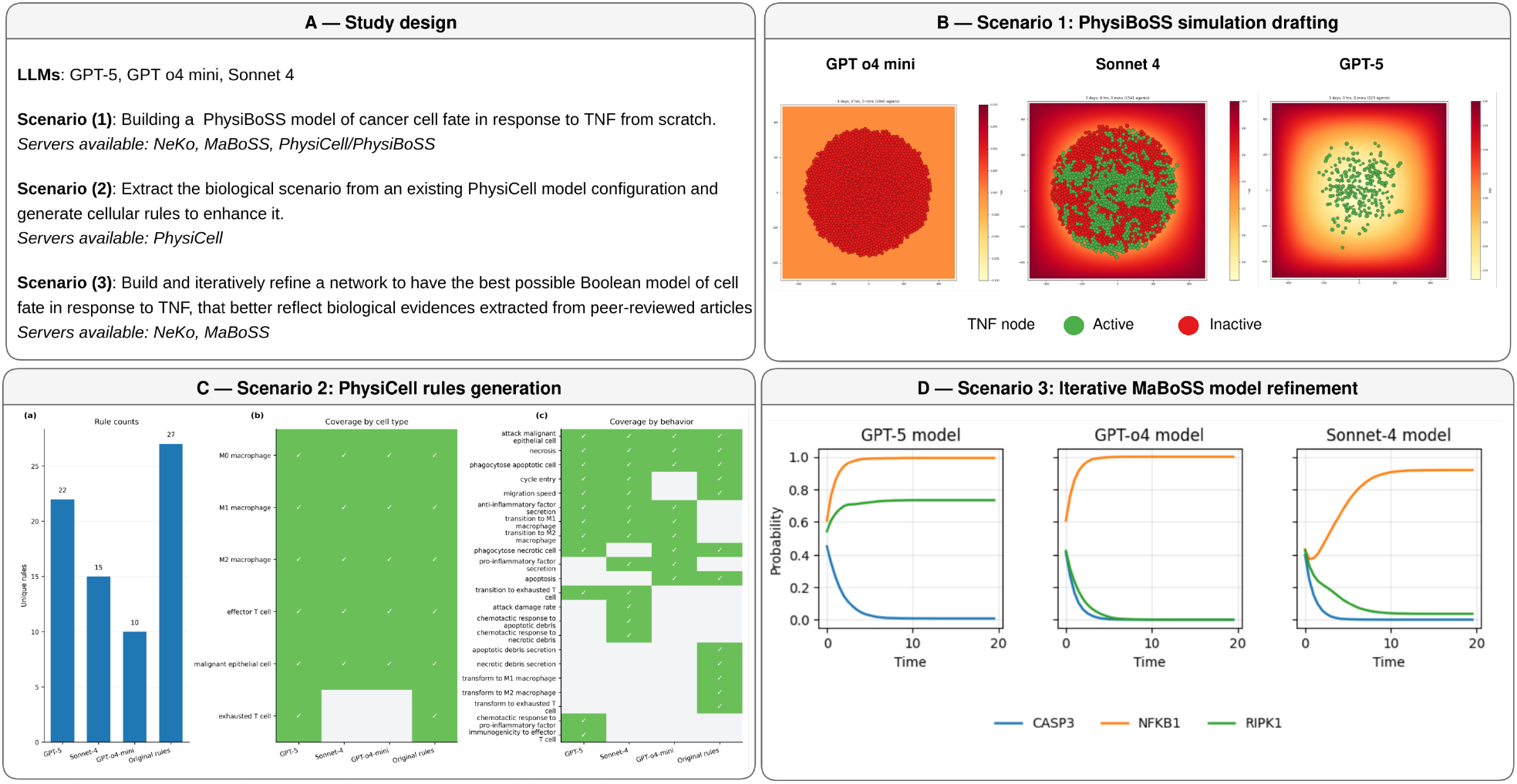
Cross-LLM comparative overview. Study design and scenario-wise summaries across LLMs.

### A. Scenario 1 — Building a multiscale model of cancer cell fate in response to TNF

This first scenario tested whether each model could autonomously execute the full NeKo→MaBoSS→PhysiCell (PhysiBoSS integration) work-flow to construct a full multiscale model of cancer cell fate upon TNF stimulation. We initiated the interaction with a single high-level prompt specifying: (i) the biological objective (“build a model of cancer cell fate in response to TNF capable of capturing apoptosis or proliferative outcomes”), and (ii) a coarse workflow outline: start from an initial gene pool, create a molecular interaction network with NeKo using OmniPath, export to MaBoSS for Boolean simulation to identify attractors, then generate a PhysiCell configuration specifying domain size, microenvironment substrates, cell types, and MaBoSS input/output mappings to phenotype rules. No explicit gene list size, network depth, or parameter values were provided in the initial prompt to assess autonomous tool selection and parameters input. For this scenario we checked whether 1) the LLMs constructed a network that includes at least some pivotal genes involved in TNF response (such as TNF receptors, Caspases for death signaling and NFkB for proliferative response), 2) generated a functioning MaBoSS model selecting valid output nodes, 3) included diffusible TNF substrate and cancer cell type in the PhysiCell configuration, with proper output/input node mapping for MaBoSS. All three LLMs completed the required tool chain with only occasional confirmation for executing the tasks. The generated multiscale models diverged along three main axes: Boolean network size/complexity, MaBoSS attractor structure, and richness of PhysiCell parameterization and MaBoSS–PhysiCell coupling. It is important to mention that we did not request to the LLMs to tune the MaBoSS model for reaching the best possible attractors.

*GPT o4 mini* produced a Boolean network with ∼50 nodes and ∼320 interactions with a minimal PhysiCell configuration, containing a cancer cell type and TNF as a substrate but with missing diffusion / uptake parameter specification, leading to absent TNF gradient formation and no activation of designated input nodes (Fig 3 panel B, GPT o4 model). The dominant MaBoSS attractor corresponded to a proliferative / survival state, consistent with insufficient extracellular pro-apoptotic stimulus.

*Sonnet 4* generated a larger and more connected network (∼200 nodes, >2,500 interactions prior to pruning bimodal / duplicated edges) and achieved a more balanced attractor landscape (quiescence vs. cycling) in MaBoSS simulations. Its PhysiCell configuration specified 3D domain geometry, oxygen and TNF diffusion/decay coefficients, secretion/uptake parameters, and mapped MaBoSS outputs to cell cycle progression and death pathways, enabling dynamic TNF exposure of cells (Fig 3 panel B, Sonnet 4 model). Despite the more advanced configuration, no cell death was triggered due to the absence of apoptotic signals in the attractor landscape.

*GPT-5* converged on a network with the smallest size compared to the other LLMs (34 nodes and 170 interactions) while providing the most explicit parameterization: boundary (Dirichlet) TNF conditions (e.g. TNF diffusing at constant rate from the border of the domain), oxygen diffusion, apoptotic and necrotic rate constants, and threshold mappings between MaBoSS nodes (e.g., NF-κB, CASP3) and Physi-Cell signals/behaviors. This leaded to the correct activation of MaBoSS input nodes in response to microenvironmental TNF and appropriate triggering of death vs. survival behaviors.

All generated configuration files were correctly executed in PhysiCell-Studio with minimal adjustment (we restricted the simulation to 2D to avoid long computation time), indicating proper tool orchestration despite the absence of details in the initial prompt.

Limitations of this scenario include the absence of systematic literature cross-validation within the conversational loop and lack of automated parameter fitting.

We have reported the full chat with all the agents in the Supplementary Material at https://github.com/marcorusc/Supp_mat_MCP_orchestrator.

### B. Scenario 2 — Rule extraction/extension for an existing PhysiCell configuration

As previously mentioned, PhysiCell allows the specification of cellular dynamics through a natural language interpretation. We tested the capability of different LLMs to extract biological information from an existing PhysiCell configuration file (https://github.com/MathCancer/PhysiCell/blob/master/sample_projects/rules_sample/config/PhysiCell_settings.xml) and to generate rules to enrich the simulation. This scenario was meant to challenge the LLM into combining the tools from a single server (PhysiCell) with its own ability of extracting and summarizing information (thus composing the biological scenario). The rules generated by each model were then compared against the original, manually curated ones. Given the input XML, all models were able to infer the biological scenario and synthesize plausible rules. *GPT-4 mini* generated 10 rules, spanning almost all of the defined cell types except exhausted T cells, and did not provide explicit Hill parameters. *Sonnet-4* generated 15 rules, covering the same cell types as GPT-4 mini. *GPT-5* produced 22 rules, including updates in CSV/XML format, and covered all cell types defined in the configuration file (Fig. 3, panel C). Compared to the manually curated rules, which included 27 entries, the LLMs proposed fewer rules overall but introduced a broader repertoire of behaviors. The curated rules emphasized core processes such as apoptotic and necrotic debris secretion—signals that are central for immune cell motility—whereas these dynamics were inconsistently captured by the LLMs. Nevertheless, the three LLMs converged on highly similar behaviors, with *GPT-5* and *Sonnet-4* additionally suggesting dynamics absent from the original model.

### C. Scenario 3 — Iterative Boolean model refinement through network pruning

The third scenario explored whether LLMs can identify flaws or inconsistencies in automatically generated Boolean models and improve their dynamics through iterative refinement, by solely using the tools from the NeKo and MaBoSS servers. We prompted the LLMs to create a workflow of the form *build–prune–convert–simulate–mutate–curate*. In practice, the LLMs first constructed a regulatory network with NeKo, autonomously detected inconsistent pathways, pruned them, converted the network into a MaBoSS model, simulated the wild-type (WT) dynamics (e.g. random initial conditions), and finally tested a set of mutations to assess biological plausibility. Importantly, the initial prompt contained no explicit guidance about which genes to chose as output or which mutants to test, in order to maximize the reasoning and orchestration performed by the LLMs themselves.

All models successfully executed the workflow, differing mainly in the number of refinement iterations. *GPT-4 mini* iterated the least, whereas *Sonnet-4* and *GPT-5* performed multiple cycles before providing their final outputs. Despite these procedural differences the resulting WT simulations were broadly consistent. We have selected 3 nodes as proxy to monitor proliferative (*NFKB*), apoptotic (*CASP3*) and necrotic (*RIPK1*) states across the generated network (22, 23). All three models displayed high levels of *NFKB1* (24), a transcription factor generally associated with cell survival, and low levels of *CASP3*, a pro-apoptotic gene linked to cell death. This mutual exclusivity between survival and death markers has also been reported in manually curated cancer cell fate Boolean models (22), suggesting that the automatically generated models can capture at least some expected dynamics (Fig 3 panel D).

A key difference emerged in the regulation of *RIPK1*. In the models generated by *Sonnet-4* and *GPT-4 mini, RIPK1* remained inactive throughout the simulations, whereas in the model generated by *GPT-5* it displayed a clear activation pattern. This divergence likely reflects differences in how each LLM orchestrated pruning and model’s parameterization during the refinement process, rather than a direct biological conclusion. A full report of the simulations and tested mutations is available in the Supplementary Material.

### D. Final considerations

MCP servers enable LLM-agnostic orchestration, but the choice of the model still influences the resulting networks in terms of size, logic depth, and emergent attractors. We found that prompt design plays a central role: increasing specificity, exposing tool calls, and clearly defining boundary conditions (e.g., TNF diffusion sources), output nodes, and mutation panels can help reduce bias toward single-attractor outcomes and improve interpretability. In this study we deliberately focused on a one-shot orchestration mode, which demonstrated that LLMs can autonomously manage complex workflows. Nevertheless, better results may be obtained by adopting a conversational mode, where tool calls are refined iteratively in dialogue with the user (see Fig 2, panel B).

## AI-Assisted Rapid Prototyping: Capabilities and Current Limitations

### A. Rapid Prototyping and Biological Reasoning

AI-assisted mechanistic modeling fundamentally transforms the time scales of computational hypothesis testing. Traditional model development requires months of literature research, coding, parameter fitting, and expert consultation. Our approach compresses this initial prototyping phase to minutes while maintaining biological logic, allowing rapid exploration of multiple hypotheses and scenarios. This acceleration lowers the barrier between biological questions and computational exploration, making computational modeling accessible to a broader community.

Importantly, the generated models demonstrate more than pattern matching. The ability of LLMs to integrate across molecular, cellular, and tissue levels suggests genuine multiscale biological competence. When properly coupled with domain-specific tools, AI assistants can function as knowledgeable laboratory partners rather than simple tool orchestrators. This capability scales with model sophistication: advanced LLMs manage general prompts effectively, while simpler ones require more detailed guidance.

### B. Technical and Practical Limitations

In our experience, the main bottlenecks arise from the underlying tools rather than the AI itself. Large network searches in NeKo or large Boolean models with many output nodes in MaBoSS may demand significant computational time. Failures usually stem from insufficient tool documentation or software bugs rather than flawed reasoning by the LLM. When biological choices appear suboptimal, they usually reflect general prompts requiring more specific guidance rather than fundamental reasoning failures. More detailed conversational interactions consistently improve model quality and biological specificity.

### C. Validation and Reproducibility Challenges

Model validation remains the primary limitation of current implementations. Systematic comparison against experimental data and established models still requires manual intervention. Quality assessment of AI-generated models should combine three complementary criteria: (1) compliance with available experimental data, (2) validation by human biological experts, and (3) evaluation by specialized AI agents with access to literature databases. Our current framework lacks such protocols, but the architecture supports their integration. Reproducibility across different prompts and users also requires systematic evaluation. Preliminary observations suggest that similar biological questions yield consistent model structures, but formal studies on prompt sensitivity and user variability are still needed for widespread adoption.

## Future Directions and Community Roadmap

### A. Technical Development Priorities

This first implementation demonstrates the promise of conversational mechanistic modeling, but its evolution and consolidation will require advances across multiple technical domains. Enhanced model quality will depend on improved automated curation algorithms, sophisticated parameter optimization workflows, and robust validation protocols. Integration with experimental data streams could enable real-time model refinement and validation against emerging datasets.

Building on our server-centered ecosystem, scaling challenges must be addressed to handle genome-wide networks, detailed kinetic models, and complex multiscale systems. This includes both computational efficiency improvements and more sophisticated AI-tool integration patterns. The development of specialized LLMs trained on systems biology knowledge could improve biological reasoning and model quality, particularly for complex cross-server integration scenarios (25).

Finally, reproducibility and provenance tracking represent critical infrastructure needs. Conversational workflows must maintain the same standards of scientific reproducibility as traditional computational approaches. This requires enhanced logging, version control, and standardized reporting mechanisms that capture the full conversational modeling process while preserving intelligent tool chain discovery capabilities. At the same time, reproducibility in this setting is complicated by the inherent variability of LLMs: different prompts or sampling settings can lead to divergent outputs, and these dependencies are not yet systematically controlled. Addressing this will require not only technical solutions but also governance frameworks and guardrails that constrain tool use, record decision paths, and ensure that results remain verifiable and interpretable across users and models.

### B. Broader Implications for Scientific Practice

AI-assisted mechanistic modeling represents part of a broader transformation in scientific computing toward agentic systems development. Recent work on multi-agent scientific problem solving (26) demonstrates how specialized AI agents can collaborate to solve complex problems across domains. Our MCP servers provide foundational orchestration capabilities that enable such collaborative approaches in systems biology.

The long-term vision encompasses integrated agent ecosystems where specialized AI assistants collaborate on model construction, validation, and optimization. A model-building agent (demonstrated here) would collaborate with literature synthesis agents, parameter optimization specialists, and experimental data integration systems. Each agent would communicate through natural language and structured file exchange, creating human-readable workflows that maintain scientific transparency.

Such systems could reshape collaboration patterns by enabling faster hypothesis-to-experiment cycles and democratizing access to computational tools. This shift may increasingly focus researchers on higher-level conceptual work, hypothesis generation, and biological interpretation, while AI systems handle routine computational tasks.

## Conclusions

Intelligent tool orchestration through MCP servers represents a promising advancement in AI-assisted mechanistic modeling for systems biology. Our example of TNF response modelling developed here, demonstrates how AI assistants can function as knowledgeable laboratory partners, exhibiting sophisticated biological reasoning while rapidly prototyping complex multiscale models through natural language interactions.

In this work we presented the following key contributions: 1) we proposed an architectural framework for biological AI–tool integration, in which modeling tasks are decomposed into decision-based tools and coordinated through session-aware orchestration. 2) we showed that modern LLMs can exhibit biological reasoning capabilities when coupled to specialized tools. 3) we provided a proof-of-concept for rapid model prototyping that reduces traditional development timelines while preserving biological logic. These elements are not only expected features of effective AI-assisted modeling but also represent our proposed framework for how such systems can be implemented in computational biology.

However, realizing this potential requires development of validation protocols, quality control mechanisms, and further integration with established computational biology framework (such as Compucell3D, Biodynamo, Corneto, CellNopt, etc.).

The developments presented here — and others that are surely underway (14, 26) — are only the beginning of a broader vision. We anticipate the emergence of multi-agent collaborative systems, in which specialized AI assistants coordinate on model construction, validation, and optimization. Realizing this vision will demand not only human oversight, but will require the appropriate tools and systems to make it effective. Maintaining scientific rigor must be a priority, requiring appropriate control mechanisms and systematic grounding of all developments in biological evidence.

Our implementation supports human control through session-aware orchestration, which makes tool calls transparent and traceable, while structured outputs and informative error handling provide interpretable checkpoints where users can intervene. Scientific rigor is reinforced by relying on established modeling software for network construction, Boolean models simulation, and multicellular dynamics, ensuring that all computations are grounded on validated methods.

The path forward requires collaboration between AI researchers, systems biologists, and the broader scientific community to ensure that these capabilities serve scientific discovery goals while addressing challenges of validation, reproducibility, and quality control in AI-assisted research. As these technologies mature, intelligent tool orchestration could become an integral component of the scientific toolkit, accelerating discovery through AI partnership while preserving the biological insight and scientific rigor essential for advancing our understanding of complex biological systems.

Although our demonstrations focus on systems biology, the principles developed here are not restricted to this domain. The same architectural ideas — modular tool orchestration, transparent workflows, and conversational interfaces — are broadly applicable to other scientific fields that depend on complex computational pipelines and heterogeneous data integration.

## Materials and Methods

We provide the whole code of the MCP servers at https://github.com/marcorusc/mcp-biomodelling-servers. The Supplementary Material containing additional analysis and the reports with the full conversations with the 3 LLMs that produced the results displayed in the article is available at https://github.com/marcorusc/Supp_mat_MCP_orchestrator. The MCP servers have been installed and executed in Visual Studio Code.

## Supporting information

Sonnet 4 chat (prompt + final results)

GPT-5 chat (prompt + final results)

GPT-o4_mini chat (prompt + final results)

Scenario analysis notebook

Links to code of the MCP servers

## ACKNOWLEDGEMENTS

We thank the developers of NeKo, MaBoSS, and PhysiCell for creating the foundational tools that made this work possible, Dr. Laurence Calzone and Dr. Eliott Jacopin for the useful comments and feedback.

## AUTHOR CONTRIBUTIONS

M.V. and M.R. conceived the project, M.R. developed the MCP servers, performed the modeling demonstration, and wrote the manuscript, A.V. supervised and funded the project.

## Bibliography

1. Chenhui Ma and Evren Gurkan-Cavusoglu. A comprehensive review of computational cell cycle models in guiding cancer treatment strategies. NPJ Systems Biology and Applications, 10:71, July 2024. ISSN 2056-7189. doi: 10.1038/s41540-024-00397-7.

2. Giulia Callegaro, Johannes P. Schimming, Janet Piñero González, Steven J. Kunnen, Lukas Wijaya, Panuwat Trairatphisan, Linda van den Berk, Kim Beetsma, Laura I. Furlong, Jeffrey J. Sutherland, Jennifer Mollon, James L. Stevens, and Bob van de Water. Identifying multiscale translational safety biomarkers using a network-based systems approach. iScience, 26(3), March 2023. ISSN 2589-0042. doi: 10.1016/j.isci.2023.106094. Publisher: Elsevier.

3. Martin Meier-Schellersheim, Iain D. C. Fraser, and Frederick Klauschen. Multiscale modeling for biologists. WIREs Systems Biology and Medicine, 1 (1):4–14, 2009. ISSN 1939-005X. doi: 10.1002/wsbm.33. _eprint: https://onlinelibrary.wiley.com/doi/pdf/10.1002/wsbm.33.

4. Ahmadreza Ghaffarizadeh, Randy Heiland, Samuel H. Friedman, Shannon M. Mumenthaler, and Paul Macklin. PhysiCell: An open source physics-based cell simulator for 3-D multicellular systems. PLOS Computational Biology, 14(2):e1005991, February 2018. ISSN 1553-7358. doi: 10.1371/journal.pcbi.1005991. Publisher: Public Library of Science.

5. Miguel Ponce-de Leon, Arnau Montagud, Vincent Noël, Annika Meert, Gerard Pradas, Emmanuel Barillot, Laurence Calzone, and Alfonso Valencia. PhysiBoSS 2.0: a sustainable integration of stochastic Boolean and agent-based modelling frameworks. npj Systems Biology and Applications, 9(1):1–12, October 2023. ISSN 2056-7189. doi: 10.1038/s41540-023-00314-4. Number: 1 Publisher: Nature Publishing Group.

6. Maciej H. Swat, Gilberto L. Thomas, Julio M. Belmonte, Abbas Shirinifard, Dimitrij Hmeljak, and James A. Glazier. Multi-Scale Modeling of Tissues Using CompuCell3D. Methods in cell biology, 110:325–366, 2012. ISSN 0091-679X. doi: 10.1016/B978-0-12-388403-9.00013-8.

7. Lukas Breitwieser, Ahmad Hesam, Jean de Montigny, Vasileios Vavourakis, Alexandros Iosif, Jack Jennings, Marcus Kaiser, Marco Manca, Alberto Di Meglio, Zaid Al-Ars, Fons Rademakers, Onur Mutlu, and Roman Bauer. BioDynaMo: a modular platform for high-performance agent-based simulation. Bioinformatics, 38(2):453–460, January 2022. ISSN 1367-4803. doi: 10.1093/bioinformatics/btab649.

8. Jörn Starruß, Walter de Back, Lutz Brusch, and Andreas Deutsch. Morpheus: a user-friendly modeling environment for multiscale and multicellular systems biology. Bioinformatics, 30(9):1331–1332, May 2014. ISSN 1367-4803. doi: 10.1093/bioinformatics/btt772.

9. Dongkwan Shin, Jeong-Ryeol Gong, Seoyoon D. Jeong, Youngwon Cho, Hwang-Phill Kim, Tae-You Kim, and Kwang-Hyun Cho. Attractor Landscape Analysis Reveals a Reversion Switch in the Transition of Colorectal Tumorigenesis. Advanced Science, 12(8):2412503, January 2025. ISSN 2198-3844. doi: 10.1002/advs.202412503.

10. Vincent Noël, Aurélien Naldi, Laurence Calzone, Loic Paulevé, and Denis Thieffry. Reproducible Boolean model analyses and simulations with the CoLoMoTo software suite: a tutorial. Interface Focus, 15(3):20250002, August 2025. doi: 10.1098/rsfs.2025.0002. Publisher: Royal Society.

11. Randy Heiland, Daniel Bergman, Blair Lyons, Grant Waldow, Julie Cass, Heber Lima da Rocha, Marco Ruscone, Vincent Noël, Paul Macklin, Daniel Bergman, Blair Lyons, Grant Waldow, Julie Cass, Heber Lima da Rocha, Marco Ruscone, Vincent Noël, and Paul Macklin. PhysiCell Studio: a graphical tool to make agent-based modeling more accessible. Gigabyte, 2024:1–19, June 2024. ISSN 2709-4715. doi: 10.46471/gigabyte.128. Publisher: GigaScience Press.

12. Marco Ruscone, Andrea Checcoli, Randy Heiland, Emmanuel Barillot, Paul Macklin, Laurence Calzone, and Vincent Noël. Building multiscale models with PhysiBoSS, an agent-based modeling tool. ArXiv, page arXiv:2406.18371v1, June 2024. ISSN 2331-8422.

13. Jeanette A. I. Johnson, Daniel R. Bergman, Heber L. Rocha, David L. Zhou, Eric Cramer, Ian C. Mclean, Yoseph W. Dance, Max Booth, Zachary Nicholas, Tamara Lopez-Vidal, Atul Deshpande, Randy Heiland, Elmar Bucher, Fatemeh Shojaeian, Matthew Dunworth, André Forjaz, Michael Getz, Inês Godet, Furkan Kurtoglu, Melissa Lyman, John Metzcar, Jacob T. Mitchell, Andrew Raddatz, Jacobo Solorzano, Aneequa Sundus, Yafei Wang, David G. DeNardo, Andrew J. Ewald, Daniele M. Gilkes, Luciane T. Kagohara, Ashley L. Kiemen, Elizabeth D. Thompson, Denis Wirtz, Laura D. Wood, Pei-Hsun Wu, Neeha Zaidi, Lei Zheng, Jacquelyn W. Zimmerman, Jude M. Phillip, Elizabeth M. Jaffee, Joe W. Gray, Lisa M. Coussens, Young Hwan Chang, Laura M. Heiser, Genevieve L. Stein-O’Brien, Elana J. Fertig, and Paul Macklin. Human interpretable grammar encodes multicellular systems biology models to democratize virtual cell laboratories. Cell, 0(0), July 2025. ISSN 0092-8674, 1097-4172. doi: 10.1016/j.cell.2025.06.048. Publisher: Elsevier.

14. Shanghua Gao, Ada Fang, Yepeng Huang, Valentina Giunchiglia, Ayush Noori, Jonathan Richard Schwarz, Yasha Ektefaie, Jovana Kondic, and Marinka Zitnik. Empowering biomedical discovery with AI agents. Cell, 187(22):6125–6151, October 2024. ISSN 00928674. doi: 10.1016/j.cell.2024.09.022.

15. Timo Schick, Jane Dwivedi-Yu, Roberto Dessì, Roberta Raileanu, Maria Lomeli, Luke Zettlemoyer, Nicola Cancedda, and Thomas Scialom. Toolformer: Language Models Can Teach Themselves to Use Tools, February 2023. arXiv:2302.04761 [cs].

16. Siyu Yuan, Kaitao Song, Jiangjie Chen, Xu Tan, Yongliang Shen, Ren Kan, Dongsheng Li, and Deqing Yang. EASYTOOL: Enhancing LLM-based Agents with Concise Tool Instruction, March 2024. arXiv:2401.06201 [cs].

17. Marco Ruscone, Eirini Tsirvouli, Andrea Checcoli, Denes Turei, Emmanuel Barillot, Julio Saez-Rodriguez, Loredana Martignetti, Åsmund Flobak, and Laurence Calzone. NeKo: a tool for automatic network construction from prior knowledge, October 2024. Pages: 2024.10.14.618311 Section: New Results.

18. Gautier Stoll, Barthélémy Caron, Eric Viara, Aurélien Dugourd, Andrei Zinovyev, Aurélien Naldi, Guido Kroemer, Emmanuel Barillot, and Laurence Calzone. MaBoSS 2.0: an environment for stochastic Boolean modeling. Bioinformatics, 33(14):2226–2228, July 2017. ISSN 1367-4803. doi: 10.1093/bioinformatics/btx123.

19. Juraj Gottweis, Wei-Hung Weng, Alexander Daryin, Tao Tu, Anil Palepu, Petar Sirkovic, Artiom Myaskovsky, Felix Weissenberger, Keran Rong, Ryutaro Tanno, Khaled Saab, Dan Popovici, Jacob Blum, Fan Zhang, Katherine Chou, Avinatan Hassidim, Burak Gokturk, Amin Vahdat, Pushmeet Kohli, Yossi Matias, Andrew Carroll, Kavita Kulkarni, Nenad Tomasev, Yuan Guan, Vikram Dhillon, Eeshit Dhaval Vaishnav, Byron Lee, Tiago R. D. Costa, José R. Penadés, Gary Peltz, Yunhan Xu, Annalisa Pawlosky, Alan Karthikesalingam, and Vivek Natarajan. Towards an AI co-scientist, February 2025. arXiv:2502.18864 [cs].

20. Dénes Türei, Tamás Korcsmáros, and Julio Saez-Rodriguez. OmniPath: guidelines and gateway for literature-curated signaling pathway resources. Nature Methods, 13(12):966–967, November 2016. ISSN 1548-7105. doi: 10.1038/nmeth.4077.

21. Luana Licata, Prisca Lo Surdo, Marta Iannuccelli, Alessandro Palma, Elisa Micarelli, Livia Perfetto, Daniele Peluso, Alberto Calderone, Luisa Castagnoli, and Gianni Cesareni. SIGNOR 2.0, the SIGnaling Network Open Resource 2.0: 2019 update. Nucleic Acids Research, 48(D1):D504–D510, January 2020. ISSN 1362-4962. doi: 10.1093/nar/gkz949.

22. Laurence Calzone, Laurent Tournier, Simon Fourquet, Denis Thieffry, Boris Zhivotovsky, Emmanuel Barillot, and Andrei Zinovyev. Mathematical Modelling of Cell-Fate Decision in Response to Death Receptor Engagement. PLOS Computational Biology, 6(3):e1000702, March 2010. ISSN 1553-7358. doi: 10.1371/journal.pcbi.1000702. Publisher: Public Library of Science.

23. Kentaro Hayashi, Vincent Piras, Sho Tabata, Masaru Tomita, and Kumar Selvarajoo. A systems biology approach to suppress TNF-induced proinflammatory gene expressions. Cell Communication and Signaling : CCS, 11:84, November 2013. ISSN 1478-811X. doi: 10.1186/1478-811X-11-84.

24. Young Ju Jeong, Hoon Kyu Oh, and Hye Ryeon Choi. Methylation of the RELA Gene is Associated with Expression of NF-κB1 in Response to TNF-α in Breast Cancer. Molecules (Basel, Switzerland), 24(15):2834, August 2019. ISSN 1420-3049. doi: 10.3390/molecules24152834.

25. Ali Essam Ghareeb, Benjamin Chang, Ludovico Mitchener, Angela Yiu, Caralyn J. Szostkiewicz, Jon M. Laurent, Muhammed T. Razzak, Andrew D. White, Michaela M. Hinks, and Samuel G. Rodriques. Robin: A multi-agent system for automating scientific discovery, May 2025. arXiv:2505.13400 [cs].

26. Bo Ni and Markus J. Buehler. MechAgents: Large language model multi-agent collaborations can solve mechanics problems, generate new data, and integrate knowledge, November 2023. arXiv:2311.08166 [cs].

