## Supplementary material for "Intelligent Tool Orchestration for Rapid Mechanistic Model Prototyping: MCP Servers as AI-Biology Interfaces": GPT-5 chat (prompt + final results)

### Scenario 1: Building a model of cancer cell fate in response to TNF from scratch

#### Prompt:

You are my multiscale modelling assistant. Together we are creating a mutliscale multicellular model of cancer cell fate in response to TNF. In order to chieve this we are going to use NeKo, MaBoSS and PhysiCell to create a simulation where an initial group of cancer cell will grow while TNF diffuses in the microenvironment. When the cells sense the TNF they will activate the input nodes of their boolean network. According to the final state of the network, they will perform behaviors such as die for apoptosis or proliferte. Those are he steps to follow: 1) create a network using Omnipath that describes cancer cell fates in response to tnf. This network should contains TNF sensing receptr as well asgenes responsible for proliferation and apoptosis. Once create the network we have to remove the bimodal interactions and check the connectivity. If the network is connected, export it in bnet file. After that generate the maboss file (.bnd and .cfg), create a simulation and select some nodes to identify the output, to represent apoptotic state and proliferative state. Once set those nodes as output, we run the maboss simulation, get the results and study the outcome (does the network make sense? are the states biologically feasible?). This is the first part. In the second part we will build the PhysiCell simulation and add maboss to the cancer cell type. Is everything clear?

#### Final results:

Here’s what I changed and exported.

TNF diffusion from edges

- TNF substrate: diffusion_coefficient=10000 μm²/min (slower), decay_rate=0.01 1/min, initial=0.
- Edge source: Dirichlet boundary enabled, boundary value=1.0 (TNF enters from domain edges and diffuses inward).
- Cancer cells: TNF uptake_rate=0.01 1/min; no secretion.

PhysiBoSS coupling (unchanged)

- Input link: TNF signal → boolean node TNF (threshold=0.1).
- Output links: CASP3 → apoptosis_rate (base 1e-4, active 0.02); E2F1 → proliferation_rate (base 0.001, active 0.02).
- Intracellular: dt=6 min, scaling=1, inheritance_global=false.

Domain and run

- Size: 1000 x 1000 x 20 µm, dx=20; 2D slab; duration: 24 h.
- Cell type: cancer_cell (Ki67_basic) with default volumes/motility; PhysiBoSS model attached (output.bnd/output.cfg).

Exports and files

- MaBoSS: output.bnd, output.cfg (already in workspace).
- PhysiCell XML: PhysiCell_settings.xml (vscode-file://vscode-app/usr/share/code/resources/app/out/vs/code/electron-browser/workbench/workbench.html) (exported at project root). Use this file with your simulator; if you want it in gpt_5/Scenario_1, I can place a copy there.

Global summary of what we achieved

- Built an OmniPath TNF-fate network, removed bimodal edges, verified connectivity, exported .bnet, generated MaBoSS .bnd/.cfg.
- Selected outputs (CASP3=apoptosis, E2F1=proliferation), ran MaBoSS; baseline under TNF favored NF-κB survival; mutations (e.g., XIAP OFF) modestly increased apoptosis.
- Created a PhysiCell session, added TNF substrate and a cancer cell type, integrated the MaBoSS model via PhysiBoSS, wired TNF signal to the network and network outputs to cell behaviors.
- Adjusted transport to model slow TNF diffusion from edges toward the center; exported an XML ready to run.

Want me to:

- Copy the exported XML into gpt_5/Scenario_1 and add a basic seeding (e.g., 200 cells in a central disk)?
- Add a gentle TNF boundary value (e.g., 0.2 instead of 1.0) or time-varying pulse?

### Scenario 2: Building a set of rules for an existing PhysiCell simulation by extracting the context directly from the configuration file

#### Prompt:

I have a PhysiCell configuration file that I would like to extend with biologically accurate rules. Load the PhysiCell configuration file, extract the biological scenario, extract the substrates, cell types and custom parameters that compose the simulation and, carefully examining the available signals and behaviors, add a series of rules to improve the model.

### Final Results:

Here's what I did and what I found.

Biological scenario

- Tumor–immune microenvironment with malignant epithelial cells, macrophages (M0/M1/M2), and T cells (effector/exhausted).
- Substrates: oxygen (Dirichlet at boundaries), apoptotic/necrotic debris (macrophage attractants), cytokines (pro- vs anti-inflammatory factors) driving polarization and T-cell function.

Extracted components

- Substrates: oxygen, apoptotic debris, necrotic debris, pro-inflammatory factor, anti-inflammatory factor.
- Cell types: malignant epithelial cell, M0 macrophage, M1 macrophage, M2 macrophage, effector T cell, exhausted T cell.
- Custom parameters:
  - `user_parameters.number_of_cells = 0`
  - `custom_data.sample` per cell type (malignant: 1.0; others: 0.0)
  - Existing ruleset reference enabled: `cell_rules.csv` at

**GitHub Copilot Response:**

Added biologically grounded rules

- Malignant epithelial cell:
  - Oxygen increases → cycle entry (half-max ~20, Hill 4)
  - Oxygen decreases → necrosis (half-max ~5, Hill 6)
  - Pro-inflammatory factor increases → immunogenicity to effector T
  - Anti-inflammatory factor decreases → immunogenicity to effector T
- M0 macrophage:
  - Pro-inflammatory factor increases → transition to M1
  - Anti-inflammatory factor increases → transition to M2
  - Apoptotic/necrotic debris increases → phagocytose apoptotic/necrotic cells
- M1 macrophage:
  - Contact with malignant epithelial cell increases → attack malignant epithelial cell
  - Pro-inflammatory factor increases → attack malignant epithelial cell and chemotaxis to pro-inflammatory factor
  - Anti-inflammatory factor decreases → attack malignant epithelial cell
- M2 macrophage:
  - Anti-inflammatory factor increases → anti-inflammatory factor secretion (positive feedback)

- Anti-inflammatory factor decreases → reduce attack on malignant epithelial cell
- Apoptotic debris increases → phagocytose apoptotic cells

- Effector T cell:
  - Contact with malignant epithelial cell increases → attack malignant epithelial cell
  - Pro-inflammatory factor increases / anti-inflammatory factor decreases → attack malignant epithelial cell
  - Pro-inflammatory factor increases → migration speed
  - Anti-inflammatory factor increases → transition to exhausted T cell
- Exhausted T cell:
  - Pro-inflammatory factor increases → transition to effector T cell
  - Anti-inflammatory factor decreases → migration speed

Exports created (in the workspace root)

- `cell_rules_gpt5_scenario2.csv` (22 rules)
- `PhysiCell_settings_gpt5_scenario2_with_rules.xml` (updated config referencing the new rules)

How to use

- Option A: Use `PhysiCell_settings_gpt5_scenario2_with_rules.xml` directly.
- Option B: Keep your original XML and point its `<cell_rules><ruleset>` to `cell_rules_gpt5_scenario2.csv` (and ensure the CSV is placed where the XML expects it), then run your simulation.

### Scenario 3: iteratevely refine a MaBoSS model through NeKo using the basic knowledge of the LLM

#### Prompt:

I want to create a Boolean model that describes how TNF affects cancer cell fate. In order to do so, I want you to create a network using Omnipath as interactions database. Once created the network, remove the bimodal interactions, export the network in bnet format and convert it into a MaBoSS model. Run the MaBoSS model (make sure to set thread count to 10 and select a subset of output nodes to reduce computation time) and get the results. If the results are biologically accurate and relevant, test some known mutations end verify the accuracy of the results. If the network is not satisfying, check important pathways within the network and individuate nodes or interactions that may not be relevant and bring inaccuracies in the model. Finally, make sure the final network always contains relevant nodes, important for our scenario. Iterate the process until the MaBoSS model is accurate enough.

### Final results:

Here's a concise summary of what was done, how the curated model differs, and why the logic changes are biologically supported.

What was achieved

- Built a TNF-driven Boolean network from Omnipath, removed bimodal edges, exported to BNET, converted to MaBoSS.
- Iteratively curated key logical rules for NF-κB, apoptosis, and necroptosis.
- Set TNF stimulus, selected outputs [RELA, CASP3, MLKL], ran MaBoSS with thread_count=10.
- Achieved baseline consistent with TNF-induced NF-κB activation (RELA=1, CASP3=0, MLKL=0).
- Validated NF-κB loss with IKBKB OFF; explored necro/apoptosis toggling via cFLIP/IAP logic.

Key differences: original vs curated (selected rules)

- TNF
  - Original: TNF = (TNFAIP3) & !(CFLAR | PTEN)
  - Curated: TNF = TNF (hold stimulus constant)
  - Rationale: treat TNF as external input during stimulation.
- TRADD
  - Original: TRADD = TNFRSF1A & !(TNF | CFLAR)
  - Curated: TRADD = TNFRSF1A & !CFLAR
  - Rationale: TRADD should be recruitable under TNF; cFLIP can inhibit DISC assembly.
- CASP8 (DISC)
  - Original: CASP8 = (TNF | TNFRSF1A | TRADD | TRAF2 | FADD) & !(CFLAR | BIRC2 | BIRC3 | NFKB1 | RELA)
  - Curated: CASP8 = (TNFRSF1A & TRADD & FADD) & !(CFLAR | BIRC2 | BIRC3)
  - Rationale: canonical DISC requires TNFRSF1A/TRADD/FADD; cFLIP and cIAPs suppress caspase-8.
- IKK/NF-κB activation and IκBα
  - RELA
    - Original: required IKK plus upstream nodes and !CASP3 or !NFKBIA variants
    - Curated: RELA = (IKBKB & IKBKG & CHUK) & (TNF | TNFRSF1A | TRADD | TRAF2 | RIPK1)
  - NFKBIA
    - Original: NFKBIA = (TRAF2 | CASP3 | IKBKG | NFKB1) & !(TNF | TNFRSF1A | RELA | TNFAIP3)
    - Curated: NFKBIA = (RELA | NFKB1) & !(IKBKB & IKBKG & CHUK)
  - Rationale: IKK complex (IKKβ/IKBKB, NEMO/IKBKG, IKKα/CHUK) activates NF-κB; IκBα is an NF-κB target and is degraded by IKK.
- A20 (TNFAIP3)
  - Original: TNFAIP3 = (TNF | TNFRSF1A | NFKB1 | RELA)
  - Curated: TNFAIP3 = (NFKB1 | RELA)

    - Rationale: A20 is an NF-κB–induced negative feedback, not a direct TNF output.
- TRAF2 and IKBKG (NEMO)
    - Original: TRAF2 inhibited by TNFAIP3; IKBKG inhibited by TNFAIP3
    - Curated: TRAF2 = (TNFRSF1A | RELA); IKBKG = (TNF | TNFRSF1A | TRADD | TRAF2 | RIPK1 | FADD | CASP8)
    - Rationale: relieve overly strong A20 gating to allow initial IKK activation under TNF.
- cFLIP (CFLAR)
    - Original: constitutively ON via negations (blocking DISC)
    - Curated: CFLAR = (NFKB1 | RELA | AKT1)
    - Rationale: cFLIP is NF-κB/AKT-inducible; initial DISC can form before feedback.
- cIAPs (BIRC2/3)
    - Original: BIRC2/3 activated broadly and also drove RIPK1
    - Curated: BIRC2 = (NFKB1 | RELA), BIRC3 = (NFKB1 | RELA)
    - Rationale: cIAPs are NF-κB targets; they stabilize complex I and suppress necrosome.
- RIPK1 and RIPK3/MLKL (necroptosis)
    - RIPK1
        - Original: (TNFRSF1A | TRADD | TRAF2 | FADD | BIRC2 | BIRC3) & !(CFLAR | CASP8 | TNFAIP3)
        - Curated: (TNFRSF1A | TRADD | TRAF2 | FADD) & !(CFLAR & CASP8)
    - RIPK3
        - Original: (MAP3K7) & !CASP8
        - Curated: RIPK3 = RIPK1 & !(CASP8 | BIRC2 | BIRC3)
    - MLKL unchanged: MLKL = RIPK3
    - Rationale: necroptosis requires RIPK1→RIPK3 when caspase-8 is inactive; cIAPs block necrosome formation; TAK1 alone should not drive RIPK3.
- Apoptosome and executioner caspases
    - CASP3
        - Original: mixed sources with strong inhibitors
        - Curated: CASP3 = (CASP8 | (APAF1 & CYCS)) & !XIAP
    - APAF1
        - Curated: APAF1 = (BAX | CYCS) & !XIAP
    - CASP9
        - Curated: CASP9 = (APAF1 | CYCS) & !BCL2L1
    - Rationale: align with intrinsic apoptosis control by XIAP and BCL2 family, and extrinsic via CASP8.

Behavior after curation (representative)

- Baseline TNF ON: RELA=1 (NF-κB active), CASP3=0, MLKL=0.
- IKBKB OFF: NF-κB off (`<nil>`=1) as expected.
- RIPK3 OFF: NF-κB remains on, necro off (expected).
- CASP8 OFF: necro not engaged unless IAP/cFLIP axis is also low (biologically consistent with requirement for cIAP loss or cFLIP down).

References supporting the curated logic

- NF-κB activation via IKK complex; IκBα negative feedback and IKK-mediated degradation:

    - Hayden & Ghosh. NF-κB in immunobiology. Cell Res 2011; Karin & Ben-Neriah. NF-κB at 20. Nat Immunol 2000; Hoffmann et al. The IκB–NF-κB signaling module. Science 2002.
- A20 (TNFAIP3) as NF-κB–induced negative regulator of TNFR signaling and RIPK1 ubiquitination:
    - Wertz et al. De-ubiquitination and A20 function. Nature 2004; Boone et al. A20 and NF-κB. Nat Immunol 2004; Shembade & Harhaj. A20 in NF-κB termination. Trends Immunol 2012.
- cFLIP as NF-κB/AKT-inducible DISC inhibitor:
    - Krueger et al. FLIP inhibits caspase-8. Curr Biol 2001; Micheau et al. DISC regulation. J Biol Chem 2002; Kataoka et al. FLIP expression and NF-κB. Mol Cell 2000.
- cIAP1/2 (BIRC2/3) promote complex I (NF-κB) and suppress RIPK1-dependent cell death:
    - Varfolomeev & Vucic. cIAPs in TNF signaling. Cell Death Differ 2011; Bertrand et al. cIAPs and RIPK1. Cell 2008.
- Caspase-8 suppresses necroptosis; RIPK1→RIPK3→MLKL drives necroptosis when caspase-8 is inhibited:
    - Oberst et al. Caspase-8 and necroptosis. Nature 2011; He et al. MLKL mediates necroptosis. Cell 2009/2013; Sun et al. MLKL requirement. Cell 2012; Degterev et al. Necrostatin. Nat Chem Biol 2005.
- Extrinsic and intrinsic apoptosis wiring:
    - Ashkenazi & Dixit. Death receptors. Science 1998; Li et al. Apaf-1 and cytochrome c. Cell 1997; Zou et al. Apaf-1. Cell 1997; Deveraux & Reed. IAPs inhibit caspases. Genes Dev 1999; Youle & Strasser. BCL-2 family. Nat Rev Mol Cell Biol 2008.

If you want, I can package the curated BNET and the MaBoSS outputs, or continue refining to make necro/apoptosis engage under appropriate combined perturbations.
