## Supplementary material for "Intelligent Tool Orchestration for Rapid Mechanistic Model Prototyping: MCP Servers as AI-Biology Interfaces": GPT-o4_mini chat (prompt + final results)

### Scenario 1: Building a model of cancer cell fate in response to TNF from scratch

#### Prompt:

You are my multiscale modelling assistant. Together we are creating a mutliscale multicellular model of cancer cell fate in response to TNF. In order to chieve this we are going to use NeKo, MaBoSS and PhysiCell to create a simulation where an initial group of cancer cell will grow while TNF diffuses in the microenvironment. When the cells sense the TNF they will activate the input nodes of their boolean network. According to the final state of the network, they will perform behaviors such as die for apoptosis or proliferte. Those are he steps to follow: 1) create a network using Omnipath that describes cancer cell fates in response to tnf. This network should contains TNF sensing receptr as well asgenes responsible for proliferation and apoptosis. Once create the network we have to remove the bimodal interactions and check the connectivity. If the network is connected, export it in bnet file. After that generate the maboss file (.bnd and .cfg), create a simulation and select some nodes to identify the output, to represent apoptotic state and proliferative state. Once set those nodes as output, we run the maboss simulation, get the results and study the outcome (does the network make sense? are the states biologically feasible?). This is the first part. In the second part we will build the PhysiCell simulation and add maboss to the cancer cell type. Is everything clear?

#### Final results:

**Multiscale Simulation Summary** Session ID: f6c3f9c5… Scenario: Tumor spheroid in a TNF-diffusive microenvironment. Cancer cells sense TNF via TNFRSF1A, run a MaBoSS Boolean network, and execute apoptosis or proliferation based on CASP3 and CCND1 outputs.

— Network construction — • Seed genes: TNF, TNFRSF1A, FADD, RIPK1, TRAF2, NFKB1, RELA, BCL2, BAX, CASP3, CASP8, CCND1, MYC • Nodes: 53 Interactions: 332 • Removed 14 bimodal interactions, confirmed full connectivity • Exported BNET: `Network_1.bnet`

— MaBoSS model — • Generated `output.bnd` + `output.cfg` • Simulation parameters: time_tick=0.5, max_time=1000, samples=10000 • Output nodes: CCND1 (proliferation), CASP3 (apoptosis)

Simulation results (final state probabilities):

1. **Baseline** • CCND1-ON = 1.00 • CASP3-ON = 0.00
2. **Mutation BCL2 = OFF** • CCND1-ON = 1.00
3. **Mutation CASP8 = ON** • CCND1-ON = 1.00
4. **Mutation BAX = ON** • CCND1-ON = 1.00

*All runs ended in the proliferative attractor.*

— PhysiCell integration — • Domain: 1000×1000×1000 µm, mesh 20 µm, duration 7200 min • Substrate added: TNF (D=1000 µm²/min, decay=0.1) • Cell type: `cancer_cell` (Ki67_basic) • PhysiBoSS: – Intracellular model attached (`output.bnd`/`.cfg`) – dt=6 min, scaling=1, stochasticity=0 – Input link: intracellular TNF → TNF activation – Output links: • CASP3 → apoptosis • CCND1 → cycle entry • Cell rules mirror Boolean outputs: – CASP3↑→ apoptosis; CASP3↓→ cycle entry – CCND1↑→ cycle entry; CCND1↓→ apoptosis

### Scenario 2: Building a set of rules for an existing PhysiCell simulation by extracting the context directly from the configuration file

#### Prompt:

I have a PhysiCell configuration file that I would like to extend with biologically accurate rules. Load the PhysiCell configuration file, extract the biological scenario, extract the substrates, cell types and custom parameters that compose the simulation and, carefully examining the available signals and behaviors, add a series of rules to improve the model.

#### Final Results:

I’ve integrated biologically informed cell-behavior rules into your configuration and exported the updated XML as `PhysiCell_settings_with_rules.xml`. It includes:

- Tumor hypoxia-induced necrosis and apoptosis
- Macrophage migration toward debris, phagocytosis, and differentiation (M0→M1/M2)
- M1/M2 secretion loops of pro-/anti-inflammatory factors
- Effector T-cell tumor attack on contact

Next, place the new XML in your PhysiCell project and run your simulation engine with it, for example:

```
./myproject PhysiCell_settings_with_rules.xml
```

### Scenario 3: iteratevely refine a MaBoSS model through NeKo using the basic knowledge of the LLM

#### Prompt:

I want to create a Boolean model that describes how TNF affects cancer cell fate. In order to do so, I want you to create a network using Omnipath as interactions database. Once created the network, remove the bimodal interactions, export the network in bnet format and convert it into a MaBoSS model. Run the MaBoSS model and het the results. If the results are biologically accurate and relevant, test some known mutations end verify the accuracy of the results. If the network is not satisfying, check important pathways within the network and individuate nodes or interactions that may not be relevant and bring inaccuracies in the model. Finally, make sure the final network always contains relevant nodes, important for our scenario. Iterate the process until the MaBoSS model is accurate enough.

#### Final results:

Here's a concise summary of our workflow and key findings:

1. Network construction• Seed genes: TNF, TNFRSF1A, TRADD, FADD, CASP8, CASP3, BCL2, NFKB1 (+ later TNFRSF10B, CFLAR, PTEN)• Database: Omnipath, only signed interactions, 2-step paths (later iterations used bias/consensus settings)• Pruned all "bimodal" edges and low-impact nodes for clarity
2. Model translation & setup• Exported to `.bnet`, converted to MaBoSS `.bnd`/`.cfg`• Thread count set to 10, 10 000 samples, max_time=1000, time_tick=0.5
3. Baseline simulation (TNF input)• Unique attractor: full activation of NFκB-driven survival nodes, no apoptosis (CASP3) or necroptosis (MLKL)
4. Single-gene perturbations• BCL2 OFF, NFKB1 OFF, CASP8 OFF, RIPK1 OFF, CFLAR OFF, XIAP OFF → no switch to apoptotic or necroptotic attractors
5. Co-stimulation & inhibitor scenarios• TNF + TRAIL (TNFRSF10B ON) → same survival attractor• XIAP OFF + TRAIL → still survival-dominant
6. Added PTEN• No impact on attractor
7. Interpretation • The network is robust: single KOs or single-ligand changes aren't enough to trigger death • Likely need combined perturbations (e.g., CFLAR+XIAP double KO) or inclusion of additional regulators (e.g., CFLAR, CASP3 co-activators)

Next steps could include systematic double-mutations or quantitative comparison of CASP3 activation probabilities against experimental values (≈0.6–0.8 under TNF+TRAIL). Let me know which scenario you'd like to explore. GitHub Copilot
