## Supplementary material for "Intelligent Tool Orchestration for Rapid Mechanistic Model Prototyping: MCP Servers as AI-Biology Interfaces": Scenario analysis notebook: Scenarios_analysis.html


In [1]:

```
import maboss
from matplotlib import pyplot as plt
import pandas as pd
import matplotlib.pyplot as plt
from wordcloud import WordCloud
from collections import Counter
import seaborn as sns
from pathlib import Path
import numpy as np
from collections import OrderedDict
import ginsim
```

#### Original PhysiBoSS TNF model¶

In [22]:

```
tnf_simple_bnd = "Boolean_network/cellfate.bnd"
tnf_simple_cfg = "Boolean_network/cellfate.cfg"
```

In [23]:

```
masim_tnf = maboss.load(tnf_simple_bnd, tnf_simple_cfg)

list(masim_tnf.network.keys())
```

Out[23]:

```
['FASL',
 'TNF',
 'TNFR',
 'FADD',
 'DISC_TNF',
 'DISC_FAS',
 'CASP8',
 'RIP1',
 'cIAP',
 'RIP1ub',
 'RIP1K',
 'IKK',
 'CASP3',
 'NFkB',
 'cFLIP',
 'BCL2',
 'BAX',
 'mROS',
 'MPT',
 'ROS',
 'ATP',
 'MOMP',
 'SMAC',
 'mcIAP',
 'Cyt_c',
 'mXIAP',
 'XIAP',
 'apoptosome',
 'NonACD',
 'Apoptosis',
 'Survival']
```

In [28]:

```
masim_tnf = maboss.load(tnf_simple_bnd, tnf_simple_cfg)

for node in list(masim_tnf.network.keys()):
    masim_tnf.network.set_istate(node, [0.5, 0.5])

masim_tnf.update_parameters(max_time=20, sample_count=10000)

masim_tnf.network.set_output(["RIP1K", "CASP3", "NFkB"])

results_tnf = masim_tnf.run()

results_tnf.plot_trajectory()
```

### Scenario 1 - MaBoSS model analysis¶

In [8]:

```
bnet_gpt5 = "gpt_5/Scenario_1/Network_1.bnet"
bnet_sonnet4 = "Sonnet4/Scenario_1/Network_1.bnet"
bnet_gpt_o4 = "gpt_o4_mini/Scenario_1/Network_1.bnet"
```

In [26]:

```
lqm = biolqm.load(bnet_gpt5)        

lrg = biolqm.to_ginsim(lqm)

edges = lrg.getEdges()
nodes = lrg.getNodeCount()
#ginsim.show(lrg)
print("GPT5 Network")
print(f"Number of nodes: {nodes}")
print(f"Number of edges: {len(edges)}")
```

```
GPT5 Network
Number of nodes: 34
Number of edges: 170
```

In [27]:

```
ginsim.show(lrg)
```

Out[27]:

In [30]:

```
import re

def count_bnet_nodes_edges(filename):
    nodes = set()
    edges = set()

    with open(filename) as f:
        header = next(f)  # skip "targets,factors"
        for line in f:
            line = line.strip()
            if not line:
                continue
            try:
                target, expr = line.split(",", 1)
            except ValueError:
                continue  # skip malformed lines

            target = target.strip()
            nodes.add(target)

            # extract regulator names from expression
            # remove operators & parentheses, keep only word tokens
            regulators = re.findall(r"[A-Za-z0-9_]+", expr)
            for r in regulators:
                if r != target:  # avoid self-edge
                    nodes.add(r)
                    edges.add((r, target))

    return len(nodes), len(edges)

# Example usage
n_nodes, n_edges = count_bnet_nodes_edges(bnet_sonnet4)
print(f"Nodes: {n_nodes}, Interactions: {n_edges}")
```

```
Nodes: 206, Interactions: 2504
```

In [24]:

```
lqm = biolqm.load(bnet_gpt_o4)        

lrg = biolqm.to_ginsim(lqm)

edges = lrg.getEdges()
nodes = lrg.getNodeCount()

print("GPT5 Network")
print(f"Number of nodes: {nodes}")
print(f"Number of edges: {len(edges)}")
```

```
GPT5 Network
Number of nodes: 53
Number of edges: 318
```

In [25]:

```
ginsim.show(lrg)
```

Out[25]:

In [17]:

```
bnd_gpt5 = "gpt_5/Scenario_1/output.bnd"
cfg_gpt5 = "gpt_5/Scenario_1/output.cfg"
bnd_sonnet4 = "Sonnet4/Scenario_1/output.bnd"
cfg_sonnet4 = "Sonnet4/Scenario_1/output.cfg"
bnd_gpt_o4 = "gpt_o4_mini/Scenario_1/output.bnd"
cfg_gpt_o4 = "gpt_o4_mini/Scenario_1/output.cfg"
```

In [18]:

```
masim_gpt5 = maboss.load(bnd_gpt5, cfg_gpt5)
masim_sonnet4 = maboss.load(bnd_sonnet4, cfg_sonnet4)
masim_gpt_o4 = maboss.load(bnd_gpt_o4, cfg_gpt_o4)
```

In [20]:

```
for node in list(masim_gpt5.network.keys()):
    masim_gpt5.network.set_istate(node, [0.5, 0.5])

for node in list(masim_sonnet4.network.keys()):
    masim_sonnet4.network.set_istate(node, [0.5, 0.5])

for node in list(masim_gpt_o4.network.keys()):
    masim_gpt_o4.network.set_istate(node, [0.5, 0.5])
```

In [24]:

```
# to add stats about network: number of nodes and number of interactions
```

In [21]:

```
list(masim_gpt5.network.keys())
```

Out[21]:

```
['FOS',
 'MYC',
 'CFLAR',
 'PTEN',
 'CCND1',
 'TNF',
 'TNFRSF1A',
 'TRADD',
 'RELA',
 'TRAF2',
 'FADD',
 'BIRC2',
 'BIRC3',
 'CASP8',
 'RIPK1',
 'TP53',
 'E2F1',
 'BCL2',
 'MAPK8',
 'AKT1',
 'MTOR',
 'CASP3',
 'MCL1',
 'BAX',
 'XIAP',
 'NFKBIA',
 'MAP3K7',
 'IKBKB',
 'JUN',
 'PIK3CA',
 'CDKN1A',
 'CDK4',
 'APAF1',
 'CASP9']
```

In [22]:

```
list(masim_sonnet4.network.keys())
```

Out[22]:

```
['CCL2',
 'NFATC2',
 'MYC',
 'EP300',
 'RB1',
 'IFNG',
 'CREBBP',
 'COMPLEX_P19838_Q04206',
 'FOS',
 'TNFAIP3',
 'GSTP1',
 'CAV1',
 'JAK2',
 'IL23A',
 'IL12B',
 'CREB1',
 'MAP2K3',
 'TRAF1',
 'MAPK11',
 'SERPINE1',
 'CCND1',
 'SP1',
 'CSF2',
 'CYLD',
 'MAPK14',
 'PTEN',
 'CFLAR',
 'APEX1',
 'STAT5A',
 'IL17A',
 'TNF',
 'TNFRSF1B',
 'FASLG',
 'IL2',
 'MAPK1',
 'TNFRSF1A',
 'SMURF2',
 'TNFRSF10B',
 'KRT18',
 'TRADD',
 'DAB2IP',
 'RELA',
 'CASP2',
 'STAT1',
 'FLNA',
 'PPP2CA',
 'TRAF2',
 'FADD',
 'FAS',
 'BIRC3',
 'TICAM1',
 'BIRC2',
 'CASP8',
 'OTUD7B',
 'RNF11',
 'RIPK1',
 'IRF1',
 'TP53',
 'RPAIN',
 'MAP3K1',
 'MAP2K7',
 'E2F1',
 'NFKB1',
 'BCL2',
 'CDK1',
 'MAPK3',
 'CYCS',
 'CASP1',
 'NFE2L2',
 'COPS6',
 'PRKCD',
 'VDR',
 'IL4',
 'CASP3',
 'ABL1',
 'PPP1CA',
 'MAP2K1',
 'ALPI',
 'BAD',
 'BCL2L1',
 'MAP2K2',
 'PRKCZ',
 'AKT3',
 'PRKACA',
 'CASP9',
 'BID',
 'TP73',
 'BAK1',
 'BBC3',
 'MAPK8',
 'BCL2L11',
 'SMPD1',
 'TP63',
 'GSK3B',
 'CTBP1',
 'AKT1',
 'BAX',
 'IL6',
 'EIF4G2',
 'ATF2',
 'MTOR',
 'STAT3',
 'CTNNB1',
 'VEGFA',
 'ATM',
 'IRS1',
 'JUN',
 'PIK3CA',
 'AKT2',
 'BIK',
 'CDKN2A',
 'BDNF',
 'CCNE1',
 'CDK2',
 'MAPK9',
 'PSEN1',
 'KAT5',
 'RAC1',
 'MAPK10',
 'CCNA2',
 'TGFB1',
 'ELAVL1',
 'COMPLEX_P20248_P24941',
 'TRAF6',
 'MAPKAPK5',
 'MDM4',
 'TRIM24',
 'FGF2',
 'CCNG1',
 'RBL1',
 'NCL',
 'MDM2',
 'MAP3K5',
 'E2F3',
 'AR',
 'PLCG1',
 'PPARG',
 'SMAD4',
 'RHOA',
 'CDKN1A',
 'PCNA',
 'DUSP1',
 'EGFR',
 'CDK3',
 'CCND3',
 'DDAH2',
 'NCOA3',
 'TERT',
 'CDC25A',
 'FOXM1',
 'CDKN1B',
 'CDK4',
 'LYN',
 'MAP2K6',
 'NGF',
 'IRAK1',
 'IL2RA',
 'CSNK2A1',
 'TGFB2',
 'MAP3K14',
 'IKBKB',
 'CHUK',
 'COMPLEX_Q09472_Q92793',
 'RPS6KA2',
 'ELP1',
 'TLR2',
 'TAB2',
 'NTRK1',
 'MAP3K8',
 'NFKBIA',
 'PAK2',
 'MEF2B',
 'PRDX3',
 'MEF2C',
 'SENP1',
 'SMAD3',
 'IL12A',
 'ROCK1',
 'MET',
 'ERBB3',
 'PGR',
 'MAP2K4',
 'HRAS',
 'RET',
 'CDC42',
 'DUSP5',
 'MAPK8IP1',
 'PPM1D',
 'CASP7',
 'EIF4E',
 'ARHGEF2',
 'CAMKK1',
 'ID1',
 'SHC1',
 'TRAF5',
 'ACTA1',
 'PTK2',
 'FGFR2',
 'PGF',
 'PXN',
 'FGR',
 'FOXO3',
 'CCL5',
 'CXCL8',
 'TSC2',
 'CBL']
```

In [23]:

```
list(masim_gpt_o4.network.keys())
```

Out[23]:

```
['TXN',
 'IFNG',
 'MYC',
 'RB1',
 'TNFAIP3',
 'MAP4K4',
 'PPARA',
 'TRAF1',
 'CCND1',
 'CFLAR',
 'MAPK14',
 'MAPK11',
 'TNF',
 'FASLG',
 'IL2',
 'TNFRSF1A',
 'TP53',
 'TNFRSF10B',
 'STAT1',
 'MAP2K7',
 'BIRC2',
 'MAP3K1',
 'FADD',
 'TRAF2',
 'CASP2',
 'HSP90AA1',
 'CASP8',
 'RIPK1',
 'LTBR',
 'DAB2IP',
 'RELA',
 'NFKB2',
 'TP53BP2',
 'PARP1',
 'PSIP1',
 'SYK',
 'NFKB1',
 'CHUK',
 'CASP3',
 'E2F2',
 'CDK6',
 'NR4A1',
 'BCL2',
 'SMPD1',
 'BAX',
 'CYCS',
 'XIAP',
 'JUN',
 'TERT',
 'E2F3',
 'PAK2',
 'CASP7',
 'BAD']
```

In [26]:

```
masim_gpt5.network.set_output(["NFKBIA", "CASP3", "BAX"])
masim_gpt5.param["max_time"]=20

masim_gpt_o4.param["max_time"]=20
masim_gpt_o4.network.set_output(["RELA", "CASP3", "BAX", "NFKB1"])

masim_sonnet4.param["max_time"]=20
masim_sonnet4.network.set_output(["RELA", "CASP3", "BAX", "NFKB1"])
```

In [27]:

```
results_gpt_o4 = masim_gpt_o4.run()
results_gpt5 = masim_gpt5.run()
results_sonnet4 = masim_sonnet4.run()
```

In [34]:

```
def new_figures_after(func):
    """Run func(); return fig objects created by it."""
    before = set(plt.get_fignums())
    func()
    after = set(plt.get_fignums())
    return [plt.figure(n) for n in sorted(after - before)]

def get_label(line):
    lab = line.get_label()
    return None if not lab or lab.startswith("_") else lab

def extract_model_data(figs):
    """Extract {label: (x, y)} from matplotlib figures."""
    data = {}
    for f in figs:
        for ax in f.axes:
            for ln in ax.get_lines():
                lab = get_label(ln)
                if lab:
                    data[lab] = (ln.get_xdata(), ln.get_ydata())
    return data

def build_color_map(*dicts):
    palette = ["#1f77b4","#ff7f0e","#2ca02c","#d62728",
               "#9467bd","#8c564b","#e377c2","#7f7f7f",
               "#bcbd22","#17becf"]
    all_labels = []
    for d in dicts:
        all_labels.extend(list(d.keys()))
    uniq_labels = list(OrderedDict.fromkeys(all_labels))
    return {lab: palette[i % len(palette)] for i, lab in enumerate(uniq_labels)}

# Generate plots (but don’t show yet)
plt.ioff()
figs_gpt5 = new_figures_after(lambda: results_gpt5.plot_node_trajectory())
figs_gpto4 = new_figures_after(lambda: results_gpt_o4.plot_node_trajectory())
figs_sonnet4 = new_figures_after(lambda: results_sonnet4.plot_node_trajectory())
figs_tnf_simple = new_figures_after(lambda: results_tnf.plot_node_trajectory())
plt.ion()

# Extract data from the generated figures
data_gpt5 = extract_model_data(figs_gpt5)
data_gpto4 = extract_model_data(figs_gpto4)
data_sonnet4 = extract_model_data(figs_sonnet4)
data_tnf_simple = extract_model_data(figs_tnf_simple)
# Consistent colors
cmap = build_color_map(data_gpt5, data_gpto4, data_sonnet4, data_tnf_simple)

# Build the 1×3 composite figure
fig, axes = plt.subplots(1, 4, figsize=(9, 3), sharey=True)
panels = [
    ("GPT-5 model", data_gpt5, axes[0]),
    ("GPT-o4 model", data_gpto4, axes[1]),
    ("Sonnet-4 model", data_sonnet4, axes[2]),
    ("Original TNF model", data_tnf_simple, axes[3])
]

for title, dct, ax in panels:
    for lab, (x, y) in dct.items():
        ax.plot(x, y, label=lab, color=cmap[lab], linewidth=1.8)
    ax.set_title(title)
    ax.set_xlabel("Time")
    ax.grid(True, alpha=0.3)

axes[0].set_ylabel("Probability")

# Shared legend below all plots
handles, labels = axes[0].get_legend_handles_labels()
unique = OrderedDict((l, h) for h, l in zip(handles, labels))
fig.legend(unique.values(), unique.keys(),
           loc="lower center", ncol=len(unique), frameon=False,
           bbox_to_anchor=(0.5, -0.15))

plt.subplots_adjust(bottom=0.25, wspace=0.3)

# Close the original generated figures
for f in figs_gpt5 + figs_gpto4 + figs_sonnet4 + figs_tnf_simple:
    plt.close(f)

plt.show()
```

### Scenario 1 - PhysiCell config file generator¶

In [90]:

```
# === PhysiCell XML macro-content (clean layout + explicit MaBoSS + cell-cycle/death summary) ===
# Columns: one per model
# Rows: (1) Domain   (2) Substrates   (3) Cell types: cycle + death   (4) MaBoSS mapping (inputs/outputs)

import xml.etree.ElementTree as ET
from pathlib import Path
import matplotlib.pyplot as plt
import matplotlib.patches as mpatches
import numpy as np
import textwrap

# ---------- INPUT ----------
XMLS = {
    "GPT-5":       "gpt_5/Scenario_1/PhysiCell_settings.xml",
    "Sonnet-4":    "Sonnet4/Scenario_1/TNF_cancer_multiscale.xml",
    "GPT-o4-mini": "gpt_o4_mini/Scenario_1/TNF_cancer_multiscale.xml",
}
MAX_SUBSTRATES     = 8      # show at most this many substrates; rest summarized
MAX_CELL_SUMMARY   = 6      # max cell types to list in the cycle/death summary
MAX_INPUT_ROWS     = 6      # max MaBoSS inputs to list
MAX_OUTPUT_ROWS    = 6      # max MaBoSS outputs to list
FIGSIZE            = (12.5, 7.0)
WRAP               = 42     # text wrap width for mapping lines

# ---------- PARSERS ----------
def _f(el, tag, default=None, cast=float):
    try:
        node = el.find(tag) if el is not None else None
        if node is None or node.text is None or node.text.strip()=="":
            return default
        return cast(node.text)
    except Exception:
        return default

def parse_xml(path):
    root = ET.parse(path).getroot()

    # Domain
    dom = root.find(".//domain")
    Dx = (_f(dom, "x_max", 0.0) - _f(dom, "x_min", 0.0)) if dom is not None else np.nan
    Dy = (_f(dom, "y_max", 0.0) - _f(dom, "y_min", 0.0)) if dom is not None else np.nan
    Dz = (_f(dom, "z_max", 0.0) - _f(dom, "z_min", 0.0)) if dom is not None else np.nan
    dx = _f(dom, "dx", np.nan); dy = _f(dom, "dy", np.nan); dz = _f(dom, "dz", np.nan)
    use2d = (dom.findtext("use_2D","false").strip().lower() in {"true","1","yes"}) if dom is not None else True
    domain = dict(use2d=use2d, Dx=Dx, Dy=Dy, Dz=Dz, dx=dx, dy=dy, dz=dz)

    # Substrates (+ Dirichlet flag)
    substrates = []
    for var in root.findall(".//microenvironment_setup//variable"):
        name = var.get("name") or var.findtext("name") or "substrate"
        d = var.find("./Dirichlet_boundary_condition")
        enabled = False
        if d is not None:
            val = (d.get("enabled") or d.text or "").strip().lower()
            enabled = val in {"true","1","yes"}
        substrates.append({"name": name, "dirichlet": enabled})

    # Cell types: cycle/death summary
    cells_summary = []
    for cd in root.findall(".//cell_definitions/cell_definition"):
        cname = cd.get("name") or cd.findtext("name") or "cell"
        cyc = cd.find(".//phenotype/cycle")
        # model name (if present) else code attribute else "-"
        model = ""
        if cyc is not None:
            model_el = cyc.find("./model")
            if model_el is not None:
                model = (model_el.get("name") or model_el.text or "").strip()
            if not model:
                model = (cyc.get("name") or cyc.get("code") or "").strip()
        if not model: model = "—"

        apop = _f(cd.find(".//phenotype/death/apoptosis"), "death_rate", None, float)
        necr = _f(cd.find(".//phenotype/death/necrosis"),  "death_rate", None, float)

        cells_summary.append({
            "name": cname,
            "cycle_model": model,
            "apoptosis_rate": apop,
            "necrosis_rate": necr
        })

    # MaBoSS mapping (explicit)
    inputs, outputs = [], []
    for cd in root.findall(".//cell_definitions/cell_definition"):
        cname = cd.get("name") or cd.findtext("name") or "cell"
        mab = cd.find(".//intracellular[@type='maboss']")
        if mab is None: 
            continue
        mapping = mab.find("./mapping")
        if mapping is None: 
            continue

        def _info(el):
            info = dict(el.attrib)
            for ch in el:
                key = ch.tag
                txt = (ch.text or "").strip()
                if txt: info[key] = txt
                for k,v in ch.attrib.items():
                    info[f"{key}.{k}"] = v
            return info

        def _fmt_params(info):
            # show at most 3 interesting params
            prefs = ["scale","weight","threshold","rate","basal","base","Kd","half_max","saturation"]
            seen = []
            for k in prefs:
                if k in info and info[k] not in ("", None):
                    seen.append(f"{k}={info[k]}")
            # add any num-like attrs if still short
            if len(seen) < 3:
                for k,v in info.items():
                    if k in {"node","name","internal_node","external_node","substrate","signal","source","target","behavior","field"}: 
                        continue
                    if isinstance(v, str) and any(ch.isdigit() for ch in v):
                        if f"{k}={v}" not in seen:
                            seen.append(f"{k}={v}")
                    if len(seen) >= 3: break
            return (", ".join(seen)) if seen else ""

        # Inputs
        for el in mapping.findall("./input"):
            info = _info(el)
            node = info.get("intracellular_name") or info.get("internal_node") or info.get("node") or info.get("name") or ""
            src  = info.get("substrate") or info.get("signal") or info.get("external_node") or info.get("source") or info.get("field") or ""
            params = _fmt_params(info)
            line = "INPUT: "
            if src:  line += f"{src} → "
            if node: line += f"{node}"
            if params: line += f"  [{params}]"
            if line != "INPUT: ":
                inputs.append(f"{cname}: {line}")

        # Outputs
        for el in mapping.findall("./output"):
            info = _info(el)
            node = info.get("intracellular_name") or info.get("internal_node") or info.get("node") or info.get("name") or ""
            tgt  = info.get("physicell_name") or info.get("behavior") or info.get("target") or info.get("field") or ""
            params = _fmt_params(info)
            line = "OUTPUT: "
            if node: line += f"{node} → "
            if tgt:  line += f"{tgt}"
            if params: line += f"  [{params}]"
            if line != "OUTPUT: ":
                outputs.append(f"{cname}: {line}")

    return {
        "domain": domain,
        "substrates": substrates,
        "cells_summary": cells_summary,
        "maboss_inputs": inputs,
        "maboss_outputs": outputs,
    }

parsed = {m: parse_xml(p) for m,p in XMLS.items()}
models = list(XMLS.keys())

# ---------- DRAW HELPERS ----------
GREEN = "#6bbf59"
LIGHT = "#f3f4f6"
BORD  = "#222"

def wrap(s, width=WRAP):
    return "\n".join(textwrap.wrap(str(s), width=width, break_long_words=False))

def draw_domain(ax, info, title):
    ax.set_title(title, pad=10)
    rect = mpatches.Rectangle((0.1, 0.5-0.30), 0.8, 0.6, fill=False, linewidth=1.1, edgecolor=BORD)
    ax.add_patch(rect)
    if info["use2d"]:
        txt = f"2D domain\nD_x × D_y = {info['Dx']:.0f} × {info['Dy']:.0f}\nΔ = ({info['dx']:.0f}, {info['dy']:.0f})"
    else:
        txt = f"3D domain\nD_x × D_y × D_z = {info['Dx']:.0f} × {info['Dy']:.0f} × {info['Dz']:.0f}\nΔ = ({info['dx']:.0f}, {info['dy']:.0f}, {info['dz']:.0f})"
    ax.text(0.5, 0.5, txt, ha="center", va="center", transform=ax.transAxes)
    ax.set_axis_off()

def draw_substrates(ax, subs):
    ax.set_title("Substrates", pad=8)
    if not subs:
        ax.text(0.5, 0.5, "None", ha="center", va="center"); ax.set_axis_off(); return
    names = [s["name"] for s in subs]
    flags = [s["dirichlet"] for s in subs]
    extra = 0
    if len(names) > MAX_SUBSTRATES:
        extra = len(names) - MAX_SUBSTRATES
        names, flags = names[:MAX_SUBSTRATES], flags[:MAX_SUBSTRATES]
    for i, (n, fl) in enumerate(reversed(list(zip(names, flags)))):  # top→bottom
        y = i
        # extend xlim a bit so right border is not clipped
        ax.barh(y, 1.0, color=GREEN if fl else LIGHT, edgecolor=BORD, height=0.8)
        ax.text(0.02, y, n, ha="left", va="center", fontsize=8)
        ax.text(0.98, y, "Dirichlet" if fl else "free", ha="right", va="center", fontsize=7, color="#444")
    if extra:
        ax.text(0.02, -0.8, f"+{extra} more…", ha="left", va="bottom", fontsize=8)
    ax.set_xlim(0, 1.03)        # <-- ensures right border shows
    ax.set_ylim(-1, len(names))
    ax.set_xticks([]); ax.set_yticks([])
    for s in ax.spines.values(): s.set_visible(False)

def draw_cell_summary(ax, rows):
    ax.set_title("Cell types — cycle & death", pad=8)
    if not rows:
        ax.text(0.5, 0.5, "None", ha="center", va="center"); ax.set_axis_off(); return
    # pick top MAX_CELL_SUMMARY (stable order)
    lines = []
    for r in rows[:MAX_CELL_SUMMARY]:
        ap = "n/a" if r["apoptosis_rate"] is None else f"{r['apoptosis_rate']:.3g}"
        ne = "n/a" if r["necrosis_rate"]  is None else f"{r['necrosis_rate']:.3g}"
        lines.append(f"{r['name']}: cycle={r['cycle_model']}; apoptosis={ap}; necrosis={ne}")
    extra = max(0, len(rows) - MAX_CELL_SUMMARY)
    y = 1.0
    step = 1.0 / (len(lines) + (1 if extra else 0) + 1)
    for ln in lines:
        ax.text(0.02, y, wrap(ln), ha="left", va="top", fontsize=8, transform=ax.transAxes)
        y -= step
    if extra:
        ax.text(0.02, y, f"… +{extra} more cell types", ha="left", va="top", fontsize=8, color="#555", transform=ax.transAxes)
    ax.set_axis_off()

def draw_maboss(ax, inputs, outputs):
    ax.set_title("MaBoSS mapping", pad=8)
    if not inputs and not outputs:
        ax.text(0.5, 0.5, "None", ha="center", va="center"); ax.set_axis_off(); return
    lines = []
    if inputs:
        lines.append("Inputs:")
        for t in inputs[:MAX_INPUT_ROWS]:
            lines.append(" • " + t)
        if len(inputs) > MAX_INPUT_ROWS:
            lines.append(f"   … +{len(inputs)-MAX_INPUT_ROWS} more")
    if outputs:
        if lines: lines.append("")  # spacer
        lines.append("Outputs:")
        for t in outputs[:MAX_OUTPUT_ROWS]:
            lines.append(" • " + t)
        if len(outputs) > MAX_OUTPUT_ROWS:
            lines.append(f"   … +{len(outputs)-MAX_OUTPUT_ROWS} more")

    y = 1.0
    step = 1.0 / (len(lines) + 1)
    for ln in lines:
        ax.text(0.02, y, wrap(ln), ha="left", va="top", fontsize=8, transform=ax.transAxes)
        y -= step
    ax.set_axis_off()

# ---------- FIGURE ----------
plt.rcParams.update({
    "font.family": "DejaVu Sans",
    "font.size": 9,
    "axes.titlesize": 10,
    "axes.labelsize": 9,
})

fig = plt.figure(figsize=FIGSIZE, constrained_layout=True)
# 4 rows × N models; tune spaces to prevent overlap
gs = fig.add_gridspec(
    nrows=4, ncols=len(models),
    height_ratios=[0.9, 1.0, 0.1, 1.8],
    hspace=0.25, wspace=0.28
)

for j, m in enumerate(models):
    dat = parsed[m]
    ax1 = fig.add_subplot(gs[0, j]); draw_domain(ax1, dat["domain"], f"{m} — Domain")
    ax2 = fig.add_subplot(gs[1, j]); draw_substrates(ax2, dat["substrates"])
    ax3 = fig.add_subplot(gs[2, j]); draw_cell_summary(ax3, dat["cells_summary"])
    ax4 = fig.add_subplot(gs[3, j]); draw_maboss(ax4, dat["maboss_inputs"], dat["maboss_outputs"])

fig.suptitle("LLM-generated PhysiCell configurations — macro-content overview", y=1.07, fontsize=11, fontweight="bold")
plt.show()

# Optional saves:
# fig.savefig("physicell_macrocontent_overview.pdf", bbox_inches="tight")
# fig.savefig("physicell_macrocontent_overview.png", dpi=600, bbox_inches="tight")
```

### Scenario 2 analysis¶

In [5]:

```
output_gpt5 = "gpt_5/Scenario_2/cell_rules_gpt5_scenario2.csv"
output_sonnet4 = "Sonnet4/Scenario_2/cell_rules_enhanced.csv"
output_gpt_o4 = "gpt_o4_mini/Scenario_2/cell_rules_with_new.csv"
original_rules = "cell_rules.csv"
```

In [16]:

```
# === Cross-LLM Rules Dashboard — light, Nature-friendly styling ===
# Panels: (a) Unique rule counts  (b) Coverage by cell type  (c) Coverage by behavior

from pathlib import Path
import pandas as pd
import numpy as np
import matplotlib.pyplot as plt
from matplotlib.colors import ListedColormap
import textwrap

# ---- INPUT ----
output_gpt5 = "gpt_5/Scenario_2/cell_rules_gpt5_scenario2.csv"
output_sonnet4 = "Sonnet4/Scenario_2/cell_rules_enhanced.csv"
output_gpt_o4 = "gpt_o4_mini/Scenario_2/cell_rules_with_new.csv"
original_rules = "cell_rules.csv"

MODEL_NAMES = {
    output_gpt5:    "GPT-5",
    output_sonnet4: "Sonnet-4",
    output_gpt_o4:  "GPT-o4-mini",
    original_rules: "Original rules"
}
HAS_HEADER       = False
TOPK_CELLTYPES   = 8     # reduce if labels still feel crowded
TOPK_BEHAVIORS   = 22
MAX_LABEL_WIDTH  = 26
SAVE_BASENAME    = "scenario2_rules_dashboard_light"

# ---- STYLE ----
plt.rcParams.update({
    "font.family": "DejaVu Sans",
    "font.size": 8.5,
    "axes.titlesize": 9.5,
    "axes.labelsize": 8.5,
    "xtick.labelsize": 8,
    "ytick.labelsize": 8,
    "legend.fontsize": 8,
    "axes.linewidth": 0.8,
    "axes.spines.top": False,
    "axes.spines.right": False,
    "xtick.direction": "out",
    "ytick.direction": "out",
    "xtick.major.size": 3,
    "ytick.major.size": 3,
    "xtick.major.width": 0.8,
    "ytick.major.width": 0.8,
    "savefig.dpi": 600,
})

# palette (closer to your original)
ACCENT  = "#1f77b4"   # bars
PRESENT = "#6bbf59"   # soft green (was darker before)
ABSENT  = "#f3f4f6"   # very light gray
CHECK   = "white"

# ---- LOAD ----
def load_rules(path: str, model_name: str, has_header=False) -> pd.DataFrame:
    df = pd.read_csv(path, header=None if not has_header else 0)
    if df.shape[1] < 4:
        raise ValueError(f"{path} has only {df.shape[1]} columns; expected ≥4.")
    df = df.iloc[:, :4].copy()
    df.columns = ["cell_type", "signal", "direction", "behavior"]
    for c in df.columns:
        df[c] = (df[c].astype(str)
                        .str.strip()
                        .str.replace(r"\s+", " ", regex=True))
    df["model"] = model_name
    df["rule_key"] = list(zip(df["cell_type"].str.lower(),
                              df["signal"].str.lower(),
                              df["direction"].str.lower(),
                              df["behavior"].str.lower()))
    df = df.drop_duplicates(subset=["rule_key"])
    return df

frames = [load_rules(Path(p), MODEL_NAMES[p], HAS_HEADER) for p in MODEL_NAMES]
data = pd.concat(frames, ignore_index=True)
models = list(MODEL_NAMES.values())

# ---- COVERAGE MATRICES ----
def presence_matrix(df: pd.DataFrame, item_col: str, topk: int) -> pd.DataFrame:
    counts = (df.groupby([item_col, "model"])["rule_key"]
                .nunique()
                .reset_index(name="n"))
    top_items = (counts.groupby(item_col)["n"].sum()
                 .sort_values(ascending=False).head(topk).index.tolist())
    counts = counts[counts[item_col].isin(top_items)]
    pres = counts.pivot(index=item_col, columns="model", values="n").fillna(0).astype(int)
    pres = (pres > 0).astype(int)
    pres["__sum"] = pres.sum(axis=1)
    pres = pres.sort_values(["__sum", *pres.columns[:-1]], ascending=[False, *([True]*len(models))])
    pres = pres.drop(columns="__sum").reindex(columns=models)
    return pres

pres_cell = presence_matrix(data, "cell_type", TOPK_CELLTYPES)
pres_beh  = presence_matrix(data, "behavior",  TOPK_BEHAVIORS)

# ---- UTILS ----
def wrap_labels(labels, width=24):
    return ["\n".join(textwrap.wrap(str(s), width=width, break_long_words=False)) for s in labels]

def panel_tag(ax, tag):
    ax.text(-0.14, 1.03, tag, transform=ax.transAxes,
            ha="left", va="bottom", fontsize=10, fontweight="bold")

# Slightly taller figure to reduce crowding
fig = plt.figure(figsize=(12.6, 6.2), constrained_layout=True)
gs  = fig.add_gridspec(nrows=1, ncols=3, width_ratios=[1.0, 1.25, 1.25])

# (a) counts
ax0 = fig.add_subplot(gs[0,0])
counts_per_model = (data.groupby("model")["rule_key"].nunique().reindex(models))
bars = ax0.bar(counts_per_model.index, counts_per_model.values, color=ACCENT, width=0.65)
ax0.set_ylabel("Unique rules")
ax0.set_title("Rule counts", pad=6)
ax0.yaxis.grid(True, color="#e5e7eb", linewidth=0.6, alpha=0.8)
ax0.set_axisbelow(True)
for t in ax0.get_xticklabels():
    t.set_rotation(15)
    t.set_ha("right")
for r in bars:
    ax0.text(r.get_x()+r.get_width()/2, r.get_height()+max(0.5, r.get_height()*0.02),
             f"{int(r.get_height())}", ha="center", va="bottom", fontsize=8)
panel_tag(ax0, "(a)")

# binary heatmap helper (green/gray + checkmarks)
def binary_heatmap(ax, pres_df: pd.DataFrame, title: str):
    ax.set_title(title, pad=6)
    cmap = ListedColormap([ABSENT, PRESENT])
    im = ax.imshow(pres_df.values, aspect="auto", cmap=cmap, vmin=0, vmax=1)
    ax.set_yticks(range(len(pres_df.index)))
    ax.set_yticklabels(wrap_labels(pres_df.index, width=MAX_LABEL_WIDTH))
    ax.set_xticks(range(len(pres_df.columns)))
    ax.set_xticklabels(pres_df.columns, rotation=0)
    # minimal box, no clutter
    for s in ("top", "right"):
        ax.spines[s].set_visible(False)
    # add checkmarks only where present
    for i in range(pres_df.shape[0]):
        for j in range(pres_df.shape[1]):
            if pres_df.iat[i, j] == 1:
                ax.text(j, i, "✓", ha="center", va="center",
                        fontsize=9, color=CHECK, fontweight="bold")

# (b) cell types
ax1 = fig.add_subplot(gs[0,1])
binary_heatmap(ax1, pres_cell, "Coverage by cell type")
panel_tag(ax1, "(b)")
for t in ax1.get_xticklabels():
    t.set_rotation(15)
    t.set_ha("right")

# (c) behaviors
ax2 = fig.add_subplot(gs[0,2])
binary_heatmap(ax2, pres_beh, "Coverage by behavior")
panel_tag(ax2, "(c)")
for t in ax2.get_xticklabels():
    t.set_rotation(15)
    t.set_ha("right")

fig.suptitle("Scenario 2 — Cross-LLM rule comparison", y=1.05, fontsize=10, fontweight="bold")
plt.show()

# Save vector + PNG
Path(f"{SAVE_BASENAME}.pdf").write_bytes(b"")  # ensure path exists on some envs
fig.savefig(f"{SAVE_BASENAME}.pdf", bbox_inches="tight")
fig.savefig(f"{SAVE_BASENAME}.png", bbox_inches="tight")
print("Saved:", f"{SAVE_BASENAME}.pdf", "and", f"{SAVE_BASENAME}.png")
```

```
Saved: scenario2_rules_dashboard_light.pdf and scenario2_rules_dashboard_light.png
```

### Scenario 3 analysis¶

In [6]:

```
output_gpt5 = "gpt_5/Scenario_3/"
output_sonnet4 = "Sonnet4/Scenario_3/"
output_gpt_o4 = "gpt_o4_mini/Scenario_3/"
```

In [13]:

```
masim_gpt5 = maboss.load(output_gpt5 + "output.bnd", output_gpt5 + "output.cfg")

masim_gpt5.network.set_output(['NFKB1', 'CASP3', 'RIPK1'])
masim_gpt5.param["max_time"]=20

masim_gpto4 = maboss.load(output_gpt_o4 + "output.bnd", output_gpt_o4 + "output.cfg")
masim_gpto4.param["max_time"]=20
masim_gpto4.network.set_output(['NFKB1', 'CASP3', 'RIPK1'])

masim_sonnet4 = maboss.load(output_sonnet4 + "output.bnd", output_sonnet4 + "output.cfg")
masim_sonnet4.param["max_time"]=20
masim_sonnet4.network.set_output(['NFKB1', 'CASP3', 'RIPK1'])
```

In [14]:

```
results_gpto4 = masim_gpto4.run()
results_gpt5 = masim_gpt5.run()
results_sonnet4 = masim_sonnet4.run()
```

In [18]:

```
results_gpto4.plot_trajectory()
```

In [19]:

```
results_gpt5.plot_trajectory()
```

In [20]:

```
results_sonnet4.plot_trajectory()
```

In [35]:

```
import matplotlib.pyplot as plt
from collections import OrderedDict

def new_figures_after(func):
    """Run func(); return fig objects created by it."""
    before = set(plt.get_fignums())
    func()
    after = set(plt.get_fignums())
    return [plt.figure(n) for n in sorted(after - before)]

def get_label(line):
    lab = line.get_label()
    return None if not lab or lab.startswith("_") else lab

def extract_model_data(figs):
    """Extract {label: (x, y)} from matplotlib figures."""
    data = {}
    for f in figs:
        for ax in f.axes:
            for ln in ax.get_lines():
                lab = get_label(ln)
                if lab:
                    data[lab] = (ln.get_xdata(), ln.get_ydata())
    return data

def build_color_map(*dicts):
    palette = ["#1f77b4","#ff7f0e","#2ca02c","#d62728",
               "#9467bd","#8c564b","#e377c2","#7f7f7f",
               "#bcbd22","#17becf"]
    all_labels = []
    for d in dicts:
        all_labels.extend(list(d.keys()))
    uniq_labels = list(OrderedDict.fromkeys(all_labels))
    return {lab: palette[i % len(palette)] for i, lab in enumerate(uniq_labels)}

# Generate plots (but don’t show yet)
plt.ioff()
figs_gpt5 = new_figures_after(lambda: results_gpt5.plot_node_trajectory())
figs_gpto4 = new_figures_after(lambda: results_gpto4.plot_node_trajectory())
figs_sonnet4 = new_figures_after(lambda: results_sonnet4.plot_node_trajectory())
#figs_tnf_simple = new_figures_after(lambda: results_tnf.plot_node_trajectory())
plt.ion()

# Extract data from the generated figures
data_gpt5 = extract_model_data(figs_gpt5)
data_gpto4 = extract_model_data(figs_gpto4)
data_sonnet4 = extract_model_data(figs_sonnet4)
#data_tnf_simple = extract_model_data(figs_tnf_simple)
# Consistent colors
cmap = build_color_map(data_gpt5, data_gpto4, data_sonnet4) #, data_tnf_simple)

# Build the 1×3 composite figure
fig, axes = plt.subplots(1, 3, figsize=(9, 3), sharey=True)
panels = [
    ("GPT-5 model", data_gpt5, axes[0]),
    ("GPT-o4 model", data_gpto4, axes[1]),
    ("Sonnet-4 model", data_sonnet4, axes[2]),
    #("Original TNF model", data_tnf_simple, axes[3])
]

for title, dct, ax in panels:
    for lab, (x, y) in dct.items():
        ax.plot(x, y, label=lab, color=cmap[lab], linewidth=1.8)
    ax.set_title(title)
    ax.set_xlabel("Time")
    ax.grid(True, alpha=0.3)

axes[0].set_ylabel("Probability")

# Shared legend below all plots
handles, labels = axes[0].get_legend_handles_labels()
unique = OrderedDict((l, h) for h, l in zip(handles, labels))
fig.legend(unique.values(), unique.keys(),
           loc="lower center", ncol=len(unique), frameon=False,
           bbox_to_anchor=(0.5, -0.15))

plt.subplots_adjust(bottom=0.25, wspace=0.3)

# Close the original generated figures
for f in figs_gpt5 + figs_gpto4 + figs_sonnet4: #+ figs_tnf_simple:
    plt.close(f)

plt.show()
```

In [ ]:
